# Stepwise reorganization of chromosome conformation and nuclear organization during stem cell differentiation

**DOI:** 10.64898/2026.09.16.752023

**Authors:** Xiangru Huo, Denis. L. Lafontaine, Thomas M. Reimonn, Liyan Yang, Richard Siller, Jack Huey, Jiang Min, Aleksandra A. Galitsyna, Vedat Yilmaz, Krishna Mohan Parsi, Emily Navarrete, Hoang Tran, Leonid Mirny, Rene Maehr, Johan H. Gibcus, Job Dekker, Nezar Abdennur

**Affiliations:** Department of Systems Biology, UMass Chan Medical School, Worcester, MA, USA; Howard Hughes Medical Institute, Chevy Chase, MD, 20815; Department of Genomics and Computational Biology, UMass Chan Medical School, Worcester, MA, USA; Program in Molecular Medicine, UMass Chan Medical School, Worcester, MA, USA; Diabetes Center of Excellence, UMass Chan Medical School, Worcester, MA, USA; Institute for Medical Engineering and Science, Massachusetts Institute of Technology, Cambridge, MA, USA; Department of Physics, Massachusetts Institute of Technology, Cambridge, MA, USA; MIT Biomicro Center, Massachusetts Institute of Technology, Cambridge, MA, USA

**Author notes:** Contributed equally (Xiangru Huo, Denis. L. Lafontaine, Thomas M. Reimonn, Liyan Yang). corresponding authors: Rene Maehr; Johan Gibcus; Job Dekker; Nezar Abdennur.

## Abstract

Nuclear organization is a fundamental feature of cell identity and cell fate determination. Compared with differentiated cells, pluripotent stem cells exhibit markedly distinct nuclear architecture at multiple levels, including chromosome folding, lamina association, histone modification landscapes and nuclear body organization. Although individual aspects of this reorganization have been characterized, how and in what order these features are remodeled as cells commit to a lineage remains poorly understood. Here, we temporally map nuclear reorganization across five stages of an *in vitro* differentiation system of human embryonic stem cells through definitive endoderm and hepatoblast intermediates into hepatocyte-like cells. By integrating Hi-C, RNA-seq, ATAC-seq, ChIP-seq, and CUT&RUN data, we establish a genome-wide framework linking structural, epigenetic, and transcriptional changes across differentiation. Combined with immunofluorescence imaging, chromosome painting, and liquid chromatin Hi-C (LC-Hi-C) we show that nuclear reorganization proceeds in a stepwise and temporally ordered manner through three major transitions. In the first transition, chromosomes condense into defined territories concurrent with transient anchoring of centromere-proximal regions to the nuclear periphery. In a second transition, active and inactive chromatin segregate, leading to stronger compartmentalization. This coincides with deposition and peripheralization of H3K9me2-marked chromatin and morphological changes in nuclear speckles, while chromatin conformation at speckle-associated regions stabilizes. In a third transition, after lineage-specific genes are activated, chromatin interactions globally stabilize, establishing a more stable nuclear architecture on top of these earlier large-scale structural rearrangements. Together, our results define a stepwise framework for nuclear reorganization during human embryonic stem cell differentiation and reveal that the transition from pluripotency to a differentiated state proceeds through coordinated, temporally ordered structural events.

## Introduction

The organization of the genome within the interphase nucleus has been extensively studied. Chromosomes form separate territories and are internally compartmentalized so that active and inactive chromatin domains of different types are spatially segregated ^1–4^. Inactive chromatin domains tend to localize near the nuclear periphery, while active domains are located towards the nuclear interior, where in some cases, they can interact with nuclear bodies such as nuclear speckles, and with (active) domains on other chromosomes. At a finer scale, chromatin loops form through the action of cohesin or through affinity-driven associations between *cis*-elements or microcompartment domains ^5–8^.

Much has been learned about how the nucleus acquires this organization by studying the process of re-building of the interphase nucleus during mitotic exit in cycling differentiated cells. Well-established cell cycle synchronization approaches have been instrumental in characterizing the order of events during this process, and to study roles of specific factors such as cohesin. As cells enter G1, the mitotic conformation is rapidly lost as condensin-mediated structures are removed during anaphase and telophase ^5,9–11^. Chromatin decondenses, and H3K27ac-marked open promoters briefly associate with each other to form a transient micro-compartmentalized state ^5–7^. Cohesin re-associates by late telophase to form loops and topologically associating domains (TADs), and over the next few hours compartments and sub-compartments appear while sub-nuclear structures such as the nucleolus, the lamina, and nuclear speckles reform.

Less understood is how nuclear architecture is remodeled over the much longer timescale of lineage commitment and terminal differentiation, a process that unfolds over days to weeks rather than minutes to hours, and that must coordinate chromosome folding with the establishment of new epigenetic landscapes, nuclear body organization, and gene expression pattern changes ^12,13^. Although pluripotent and differentiated cells share basic principles of genome organization, they differ quantitatively in several ways: Pluripotent embryonic stem (ES) cells display an unusually open chromatin state ^14^, architectural chromatin proteins bind loosely and dynamically ^15,16^, less heterochromatin is found at the nuclear periphery, chromatin is more mobile ^17^, and chromosome territories are more loosely organized ^18^. In pluripotent cells, chromatin state is distinct in composition and histone modification: active modifications dominate and heterochromatin is less defined. These differences correlate with differences in gene expression, which is initially widespread in pluripotent cells and becomes increasingly restricted during differentiation ^15,19^. Cell state transitions are further characterized by changes in which regions of the genome are active or inactive, and these changes correlate with altered sub-nuclear positioning of these loci: For instance, at the nuclear lamina specifically, genome-lamina contacts are extensively reorganized during differentiation, with DamID signal increasing at newly recruited lamina-associated domains (LADs) as cells commit to a fate ^20–22^. Large LADs, marked by repressive H3K9me2 ^23,24^ are established, lost, or repositioned in a lineage-specific manner as cells commit to a fate, a pattern now documented across multiple cell types ^20,25–27^. Similarly, repressive H3K27me3 domains broaden and spread upon differentiation, expanding beyond the sharp, promoter-restricted peaks characteristic of the pluripotent state ^28–31^. Nuclear bodies such as nucleoli, PML bodies and nuclear speckles have also been shown to reorganize during in vitro differentiation. For example, nuclear speckles become larger and fewer in number as myotubes differentiate from myoblasts ^32^. In fact, it was shown that gene activation during a cell state transition correlates with repositioning of this gene closer to nuclear speckles and that SON, a critical nuclear speckle protein, is required for expression of activated genes ^33^. Together, these reorganization features repositioning are now understood to be functionally important for establishing and maintaining cell identity and are not merely a passive consequence of it.

The temporal logic connecting changes in genome architecture, epigenetic signals, gene expression, and nuclear body positioning remains poorly resolved. Do chromosome folding, heterochromatin repositioning, histone mark deposition, and nuclear body reorganization occur simultaneously as a single coordinated event, or do they unfold as a series of temporally and possibly mechanistically separable steps?

Existing studies have largely examined individual features: a single histone mark, a nuclear landmark, or chromatin state at one or two timepoints. As a result, the order in which these different layers of nuclear architecture and maintenance are established during lineage commitment remains unknown. Moreover, prior work on chromosome folding during differentiation has often centered on individual gene loci or developmental pathways, such as the reorganization at the Oct3/4 locus during ES cell differentiation ^34^, or was limited to single-timepoint comparisons between pluripotent and committed states ^12,35^.

Here, we use a range of approaches to characterize interphase chromosome conformation, and quantitative changes in that conformation, as human embryonic stem cells (hESCs) differentiate into hepatocyte-like cells (HLCs) through five distinct cell stages. We show that interphase genome conformation in hESCs is quantitatively different from that in differentiated cells, and that the pluripotent state transforms into the differentiated state in several steps that occur at different times over many days of differentiation. In pluripotent cells, chromosomes are decondensed and diffuse, form poorly defined territories and can be extensively intermingled. Compartmentalization is weak with active and inactive chromatin not well segregated. Chromatin interactions are generally relatively short-lived. In HLCs, all these properties have changed: chromosomes are more compacted, form well-isolated territories that intermingle little, active and inactive chromatin domains are well segregated, and chromatin interactions become more stable first at speckle-associated domains, and then genome-wide. These changes coincide with large-scale changes in the epigenome, especially the deposition of heterochromatic marks and changes in the appearance and composition of nuclear landmarks including nuclear speckles and the nuclear periphery. Analysis of chromosome and nuclear organization throughout the five stages of differentiation towards HLCs uncovers a temporally ordered, stepwise framework for the dynamics of nuclear organization during embryonic stem cell differentiation, showing that distinct architecture features are established across separate time windows rather than through a simultaneous change.

## Results

### A staged in vitro hESC-to-HLC system resolves lineage commitment and its regulatory program

To resolve the temporal order in which the nucleus is remodeled during human cell-fate commitment and maturation, we generated a five-stage differentiation trajectory from H1 human embryonic stem cells (hESCs) to hepatocyte-like cells (HLCs) through a stepwise hepatic differentiation protocol (see Methods; **Fig. 1a**) extending the 4D Nucleome Consortium’s previous ESC-to-Definitive Endoderm (DE) protocol ^36^. hESCs were taken sequentially through definitive endoderm (DE; 5 days), a hepatoblast stage (HB; 8 days of hepatocyte growth factor), immature hepatocyte-like cells (iHLC; 4 days of dexamethasone) and hepatocyte-like cells (HLC; 12 days of maturation), and each stage was sampled in parallel for Hi-C, RNA-seq, ATAC-seq, and CUT&RUN/ChIP-seq for various histone modifications. Immunofluorescence for stage-specific markers confirmed the expected identity at each step (**Fig. 1b**). Principal component analysis of the transcriptomes and chromatin accessibility landscapes showed tight concordance between replicate rounds of differentiation and ordered the stages along similar differentiation axes, with the first component separating the pre-hepatic and hepatic stages, the second component separating hESC from DE, and the third component capturing deviation of HLC from HB and iHLC (**Extended Data Fig. 1a, b**).

**Fig 1:**
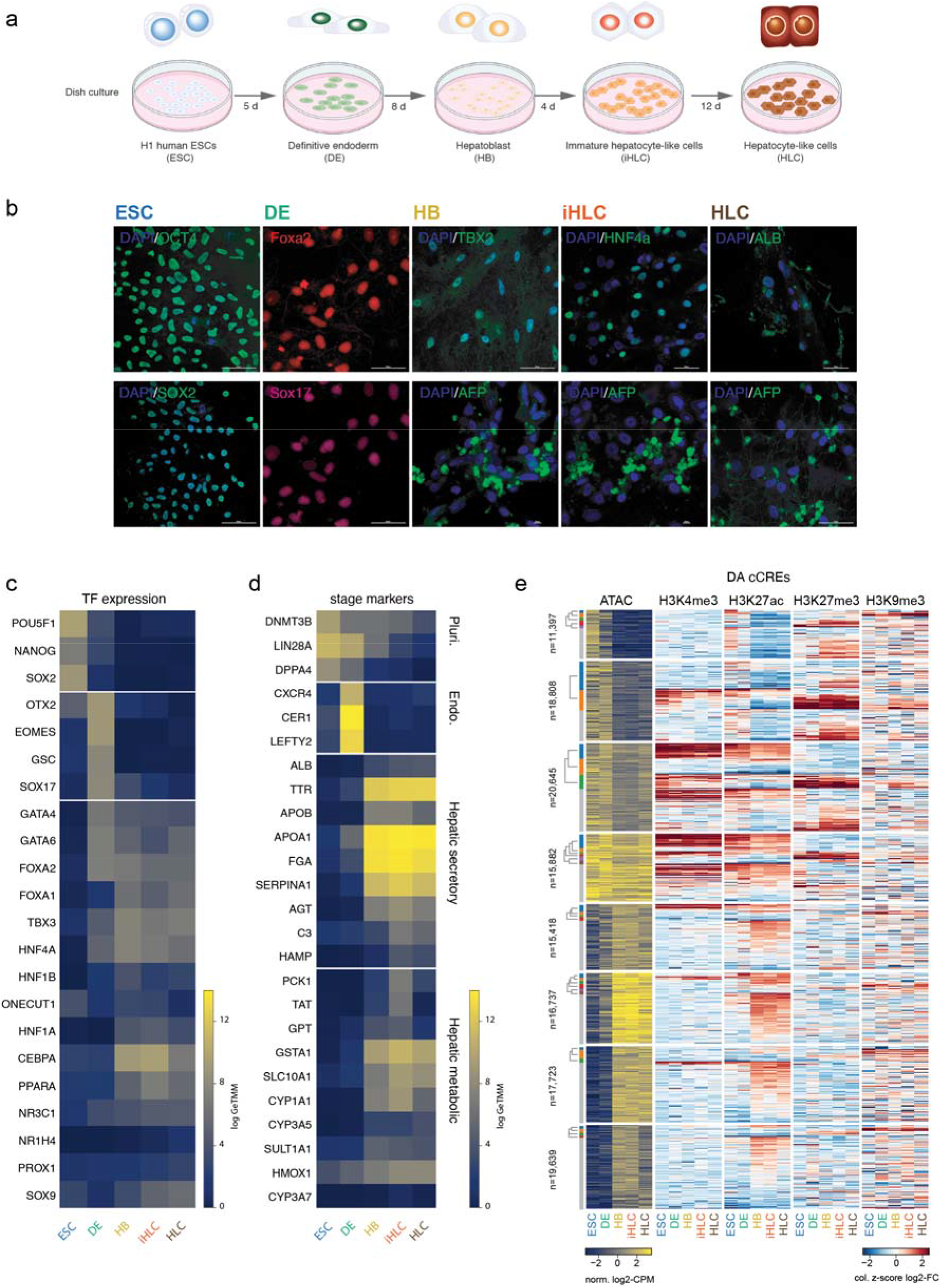
Characterization of stepwise *in vitro* differentiation trajectory of H1 human ESCs into hepatocyte-like cells. **a**, Schematic diagram of the differentiation procedure across five differentiation stages: H1 human ESCs (ESC), definitive endoderm (DE), hepatoblast (HB), immature hepatocyte-like cells (iHLC), and hepatocyte-like cells (HLC). **b**, Representative immunofluorescence images of stage-specific marker gene staining across the five stages. **c**, Expression panel for representative transcription factors across stepwise differentiation trajectory **d**, Expression panel for stage-specific marker genes. **e**, Overview of differentially accessible (DA) cCREs across the 5 stages via ATAC-seq with histone modifications. Row blocks represent fuzzy c-means clusters of accessibility trajectories of all DA cCREs. Colored bars and dendrograms on the left depict EVoC subclusters derived from the histone modification profiles on the right.

Transcriptional profiling showed that the system recapitulates the canonical, temporally ordered program of hepatic specification (**Fig. 1c**). Core pluripotency regulators (POU5F1, NANOG, SOX2) were expressed in hESCs and extinguished upon endoderm induction, which activated anterior primitive-streak and mesendoderm factors EOMES and GSC together with the definitive-endoderm specifier SOX17. The pioneer factors FOXA2 and GATA4/6 were induced as endodermal identity was acquired and maintained thereafter, accompanying the activation of the hepatoblast and hepatocyte regulators TBX3, HNF1B, HNF4A, HNF1A and CEBPA at the later stages. Functional liver markers confirmed the acquisition of hepatocyte identity (**Fig. 1d**). Secretory products characteristic of the liver, including albumin (ALB), transthyretin (TTR), and the apolipoproteins, fibrinogen and serpins (APOB, APOA1, FGA, SERPINA1), were induced from the hepatoblast stage onward, followed at the hepatocyte stages by genes of amino-acid and xenobiotic metabolism, including transaminases (TAT, GPT), glutathione S-transferases and several cytochrome P450s.

Gene-set enrichment analysis confirmed that each transition engages a coherent regulatory program (**Extended Data Fig. 1c–e**). Pathway enrichment traced the progression from proliferative pluripotency through endoderm to hepatic metabolic identity, including proliferation and cell cycle-associated gene sets among downregulated genes over the progression from hESC to iHLC, enrichment for TGF-β signaling among upregulated genes in the hESC to DE transition, and enrichment in lipid, heme, and xenobiotic metabolism gene sets among genes upregulated from DE to HB and HB to iHLC (**Extended Data Fig. 1c,d**). However, a subset of hepatoblast and terminal-maturation regulators were activated only incompletely or not at all (e.g. PROX1, ONECUT1) and many hepatic metabolic programs that peaked at iHLC were mildly reduced at HLC; meanwhile, signatures increased for focal-adhesion, proliferation, and epithelial-mesenchymal transition at the HLC stage. These observations are consistent with the documented limitations of HLC differentiation in two-dimensional culture, which fail to reach a mature hepatocyte state and may diverge to induce non-hepatic programs absent from primary hepatocytes ^37–40^. Moreover, mechanical tension arising from cell spreading in culture is sufficient to activate YAP/TAZ target genes and promote loss of hepatocyte identity, providing a plausible basis for the regression we observe over the extended maturation phase ^41^.

Chromatin accessibility was extensively remodeled across the trajectory. The union of all differentially accessible ENCODE (version 4) candidate cis-regulatory elements (cCREs) partitioned into dynamic archetypes by fuzzy c-means clustering. These archetype sites were further subclustered by H3K4me3, H3K27ac, H3K27me3, and H3K9me3 normalized signals across the 5 stages. Changes in accessibility tracked the corresponding histone-modification state: gains of H3K27ac at activated elements and loss of repressive H3K27me3 or H3K9me3, and loss of H3K27ac and gain of H3K27me3 at decommissioned ones (**Fig. 1e**). Motif enrichment within stage-specific accessible elements implicated successive members of transcription-factor families at successive transitions: for example, OTX2 followed by PITX2, and the staged deployment of GATA, FOXA and HNF factors (**Extended Data Fig. 1e**). Because related factors recognize near-identical motifs, motif enrichment alone could not reliably identify which family member acts at each stage. However, comparing the underlying elements directly, sets driving family-level enrichment at consecutive transitions were largely disjoint (**Extended Data Fig 1f**). This suggests that stage-specific family members are largely recruited to newly accessible sites, rather than re-using shared sets.

Together, these data establish a temporally resolved differentiation system in which the expected transcriptional and chromatin-accessibility programs unfold in order, providing the foundation for mapping the accompanying reorganization of genome folding and nuclear organization.

### Long-range interactions resolve into distinct chromosomal interaction trajectories that undergo large-scale remodeling at lineage commitment

We next examined how genome folding is reorganized across the trajectory using Hi-C3.0 at all five stages (**Fig. 2a**). At the level of the contact maps, hESCs and DE cells were strikingly similar: both displayed comparably low ratios of intra- to inter-chromosomal contacts and relatively weak compartmentalization. When interaction frequency was plotted as a function of genomic distance, data for all cell types displayed the expected interphase decay. All decay plots showed the characteristic shoulder at ∼100kb, which represent cohesin-mediated loops. (**Extended Data Fig. 2a**). Strikingly, at the DE-to-HB transition, the ratio of intra-to inter-chromosomal contacts increases substantially. Furthermore, the compartment checkerboard visibly intensified, reflecting sharper segregation of active from inactive chromatin. This stronger compartmentalization was sustained with little further change through the iHLC and HLC stages, as quantified by saddle plots (**Extended Data Fig. 2b**).

**Fig 2:**
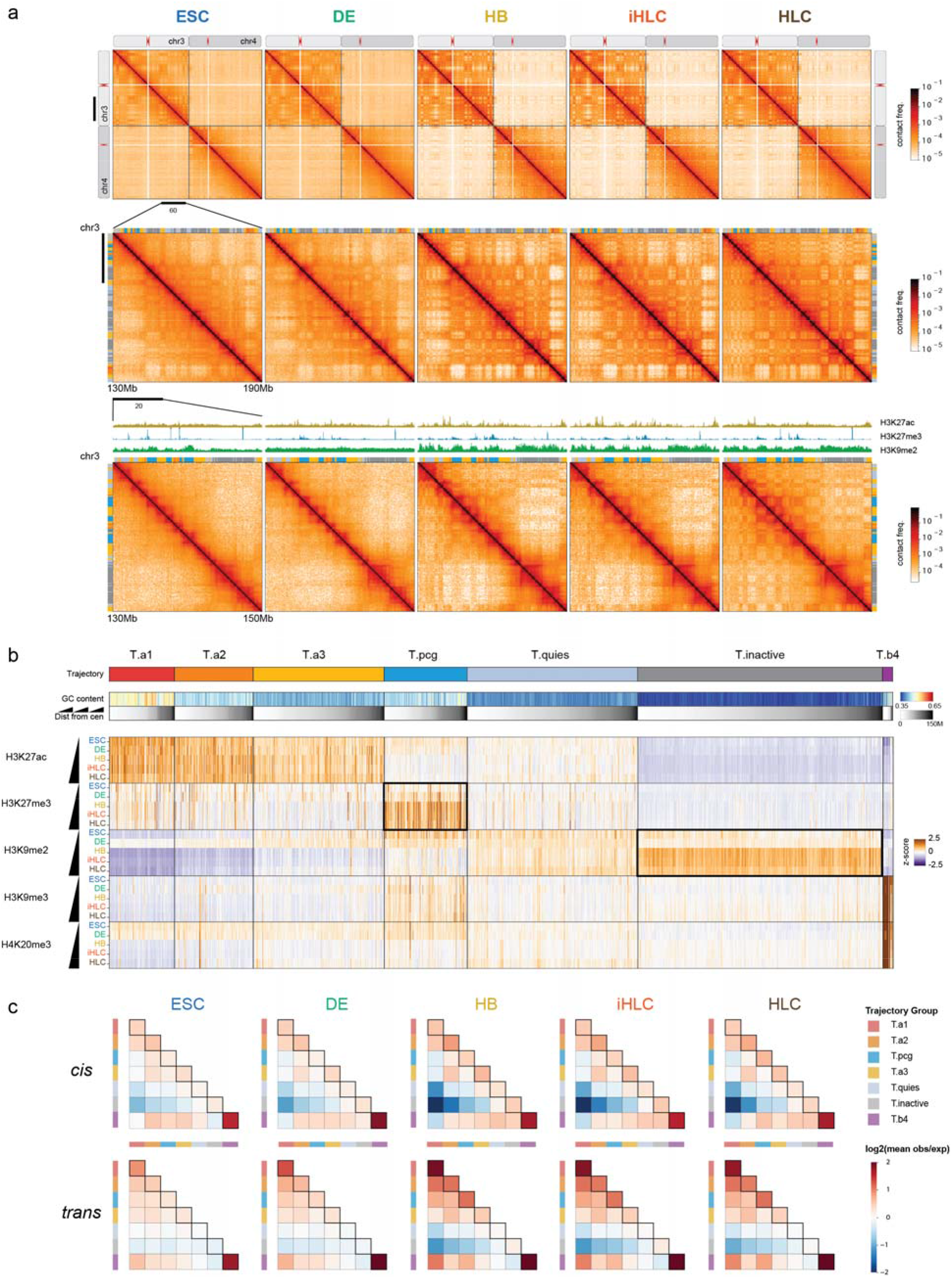
Joint analysis of long-range interactions at the locus level reveals seven broad Interaction Profile Trajectories (IPTs) across five stages. **a**, Hi-C maps of five differentiation stages (ESC to HCL) at three resolutions (whole-chromosome chr3-chr4, with zooms into ∼60-Mb then further into ∼20-Mb of a region within the chr3 q-arm) show contact frequencies alongside H3K27ac, H3K27me3 and H3K9me2 marks and IPT annotations. IPTs are denoted as the colored segment above the heatmap. The compartment checkerboard and territorial separation sharpen progressively, peaking at the HB stage and persisting thereafter. **b**, Heatmap characterizing the seven IPTs (T.a1–T.b4) consisting of bins with coherent long-range interaction dynamics across the five differentiation stages. Columns represent 50-kb genomic loci, grouped by IPT (top row, colored bars). Groups are sorted by mean GC content (second row), and columns within each group are ordered by distance from the centromere (third row). The bottom heatmap shows z-scored histone modification signal across the five stages at the same 50-kb bins. Accumulations of H3K27me3 and H3K9me2 are observed in trajectories T.pcg and T.inactive, respectively, at the DE-to-HB transition (highlighted with dark borders), while high H3K9me3 and H4K20me3 levels remain invariant within T.b4. **c**, Aggregate contact enrichment (mean observed/expected) in *cis* (top) and *trans* (bottom) between bins in IPTs depicted as heatmaps categorized by IPT at each stage of hepatic differentiation.

To localize these changes in long-range genome folding along the genome in a comparative manner, we jointly decomposed the Hi-C maps of all five stages by their long-range *trans* interaction profiles at 50-kb resolution by joint PCA ^42^. To resolve their timing, we applied K-Means clustering on the complete sequence of leading PC scores across stages for each locus. Rather than grouping loci by characteristic long-range signatures in a single sample (subcompartments / interaction profile groups), this approach seeks to capture broadly coherent patterns of long-range interactions across differentiation time, thus identifying characteristic locus trajectories ^42^. After consolidating clusters differing only in centromeric vs telomeric proximity, we identified seven distinct Interaction Profile Trajectories (IPTs) at 50-kb resolution and characterized the dynamics of their epigenetic landscapes via histone modifications, including H3K27ac, and broad repressive marks H3K27me3, H3K9me3, and H4K20me3 (**Fig. 2b. Extended Data Fig. 2c**).

Four of the seven trajectories were epigenetically stable across the time course. Three corresponded to active chromatin spanning a gradient of activity: T.a1, capturing the largely stable nuclear-speckle-associated A1 interaction profile group ^43–45^, followed by progressively less active T.a2 and T.a3. A fourth stable trajectory, which we labeled T.b4, corresponded to constitutive heterochromatin comprising pericentromeric and telomeric regions and a small number of exceptionally large H3K9me3 domains, such as those on chromosome 19 (**Fig. 2b**). Indeed, broad H3K9me3 and H4K20me3 neither spread nor changed in intensity across the differentiation. Two trajectories, labeled T.pcg and T.inactive, were dynamic with respect to repressive chromatin marks, undergoing dramatic changes as cells transition from DE to the hepatoblast stage. A final trajectory, T.quies, largely comprising regions flanking T.inactive, was quiescent in the sense of emitting little signal of any kind **(Fig. 2b)**.

The first epigenetically dynamic trajectory, T.pcg, captured domains that acquired broad, yet unevenly distributed, H3K27me3 signal at the HB stage, forming visibly distinct compartmental domains in *cis* only from HB onward (**Fig. 2a**). Its defining feature is a change in the spatial scale of H3K27me3, illustrated at for instance the Sonic Hedgehog (SHH) developmental locus (**Extended Data Fig. 3a**): in hESCs the mark formed tall, focal peaks coincident with punctate Hi-C contact dots, reminiscent of Polycomb bodies. The H3K27me3 peaks lost height and most dots in Hi-C already ceased to be visible at the DE stage. Such interaction patterns are too narrow and sparse to register by compartmental analysis at either stage. Then, from the hepatoblast stage onward, H3K27me3 instead spreads as broad, low-level domains spanning hundreds of kilobases to megabases.

This change in chromatin state had a clear transcriptional correlation. Most genes whose promoters lie in T.pcg underwent a coordinated step down in expression precisely at the DE-to-HB transition when spreading of H3K27me3 occurs, the strongest downward shift of any trajectory (**Extended Data Fig. 3b**). To find whether the silenced genes correspond to specific alternative fates rather than to generic repression, we ranked the T.pcg genes by their DE-to-HB expression change and tested for enrichment of cell-type marker sets from PanglaoDB, a broad compendium of lineage-defining genes. The down-regulated genes were significantly enriched for markers of non-hepatic lineages (predominantly neural-developmental cell types and epiblast) indicating that H3K27me deposition restricts these alternative programs as the hepatic fate is consolidated. Many of these were transiently induced at the DE stage and then collapsed at HB (**Extended Data Fig. 3b**).

To resolve where within these domains the switch enacted, we jointly clustered H3K27me3 and H3K27ac across the five stages at T.pcg gene promoters and, separately, at T.pcg cCREs (**Extended Data Fig. 3c,d**). Elements resolved into a Polycomb-poised population (high H3K27me3, low H3K27ac), an active population that gained H3K27ac, and an intermediate set that acquired H3K27me3 over the transition. Notably, in ESC, H3K27me3 was found mainly at promoters and proximal elements, which decreased as differentiation proceeded, whereas the broad H3K27me3 gained from HB onward was carried disproportionately by distal enhancers and intergenic elements, suggesting that the spreading that defines T.pcg blankets the wider regulatory domains rather than concentrating at promoters.

The second epigenetically dynamic trajectory, T.inactive, similarly gained a lasting enrichment for H3K9me2 over the same DE-to-HB window. These loci largely correspond to B compartmental regions commonly identified as lamina-associated domains by Lamin B1 DamID across 7 cell types or by SPIN states in H1 and K562 cells (**Extended Data Fig. 2d, e**). Unlike T.pcg, H3K9me2 levels rose but remained evenly distributed over the domains. Finally, we quantified compartmentalization strength, i.e., the preference for IPT loci to interact with other loci of the same IPT, across differentiation. An aggregate plot of mean observed-over-expected contact frequency between IPT loci across the five stages shows that the change in compartmentalization beginning at the DE-to-HB transition was dominated by reduced heterotypic affinity between T.a1 and T.inactive in both *cis* and *trans*, as well as increased self-affinity of T.a1 in *trans* (**Fig. 2c**).

Together, these data show that hESC differentiation is accompanied by large-scale structural and functional changes of the genome. Several major chromosomal events stand out. First, an increasing ratio of intra-to inter-chromosomal contacts suggest that ES cells have weak chromosome territories, and that during differentiation chromosome territories become more defined. Second, active and inactive chromatin domains are initially weakly segregated, but then become localized into distinct separate nuclear compartments. Third, the expansion of repressive marks H3K27me3 over alternative lineage genes and H3K9me2 across lamina-associated domains, suggests widespread establishment of repressed chromosomal domains. Below we demonstrate when each of these chromosomal events occur, where they occur within the cell nucleus and how they are associated with changes in the biophysical properties of chromatin, which led to the identification of three major temporally segregated transitions in nuclear organization that convert the pluripotent state into a differentiated state of the genome.

### Differentiation of hESCs is marked by increased chromosome territoriality and dynamic localization of centromeres and telomeres

Hi-C data shows that as cells differentiated, inter-chromosomal interactions decreased, and intra-chromosomal interactions increased (**Fig. 2a**). This suggests that chromosomes become more individualized, i.e., become more territorial. To visualize chromosome territories directly, we performed chromosome painting for chromosome 11 (chr11, 135Mb) and chromosome 15 (chr15, 107Mb) (**Fig. 3a**). In hESCs, both chromosomes occupied a relatively large nuclear volume, and the two chromosomes were often found to substantially intermingle (**Fig. 3b**). Upon exit from pluripotency, inter-chromosomal overlap decreased markedly during the transition to definitive endoderm (DE), a trend that continued throughout subsequent differentiation stages.

**Fig. 3:**
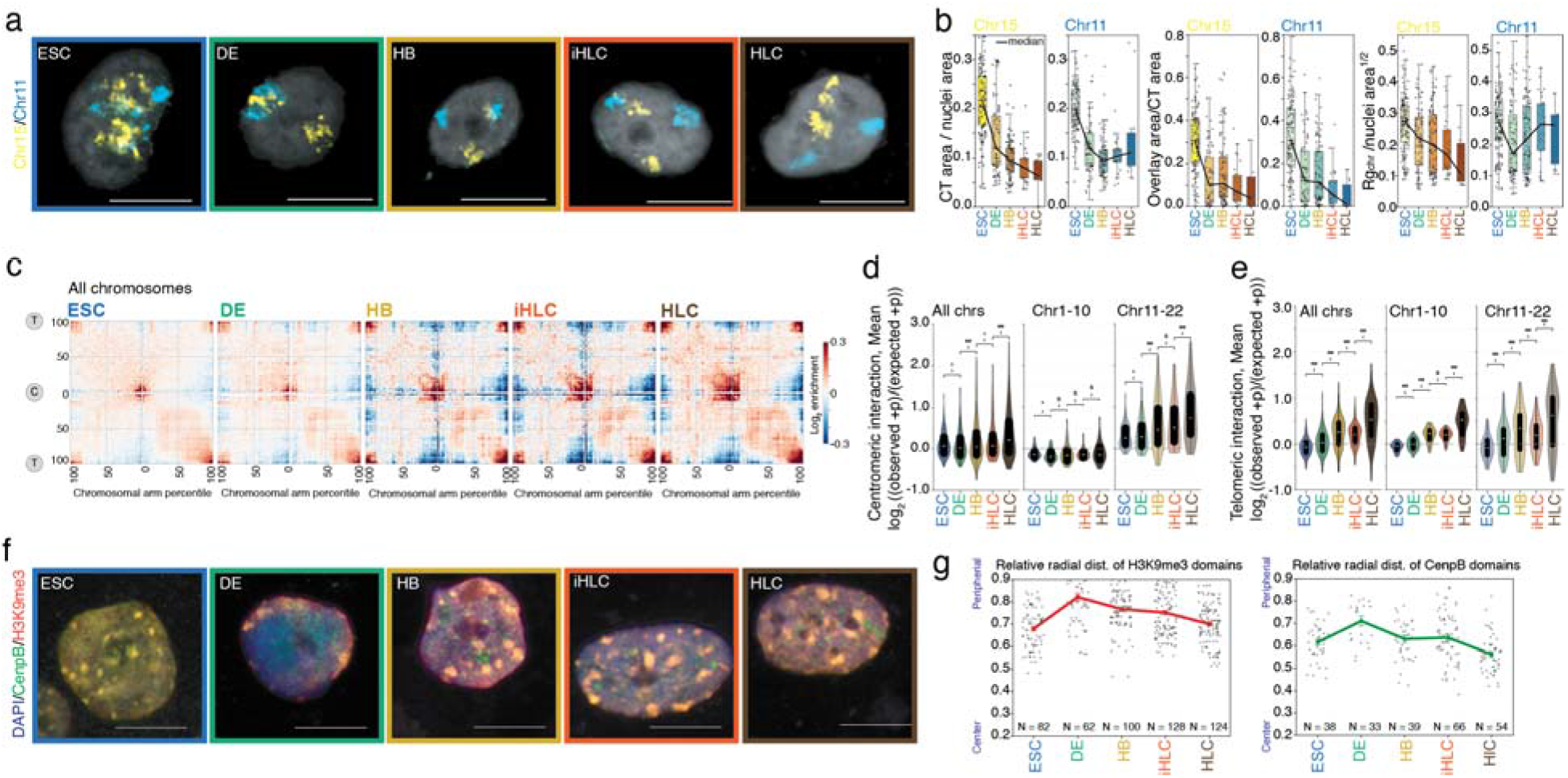
Increased chromosomal territoriality and complex dynamics of centromeres and telomeres during differentiation. **a**, A single optical section showing representative chromosome 11 (cyan) and chromosome 15 (yellow) nuclear occupancy visualized by chromosome painting across five stages. Scale bar: 10 μm. **b**, Quantifying properties of chromosome chromosomes 11 and 15 territories measured from chromosome painting across five stages. Left: ratio of chromosome territory (CT) area to total nuclear area; middle: fraction of overlapping pixels between chromosome 11 and chromosome 15 territories; right: normalized radius of gyration ((area of nucleus)^1/2^ /Radius of gyration of chromosome)) of each chromosome territory. **c**, Scaled-chromosome saddle plots showing piled-up interchromosomal interactions across all chromosomes across five stages. “T” represents telomeric regions; “C” represents centromeric regions. **d-e**, Quantifying interaction strength at pericentromeric (d) and telomeric (e) regions. Scores were calculated within 4 Mb of the corresponding telomere/centromere along each chromosome axis. Data represent the interaction strengths between different pairs of chromosomes. From left to right: all chromosomes, chromosomes 1–10 (large chromosomes), and chromosomes 11–22 (small chromosomes). *p<0.05, **p<0.01, ***p<0.001 via a Wilcoxon signed-rank test with one-sided alternative (see Methods). **f**, Representative immunofluorescence images of CenpB (green) and H3K9me3 (red) co-staining across five stages. Scale bar: 10 μm. **g**, Quantification of images in (f) showing the average relative radial distance of H3K9me3 domains (left) and CenpB domains (right) per nucleus across five stages.

Nuclear volume and chromosome compaction, quantified by the radius of gyration continuously decreased from the hESC to the hepatoblast (HB) stage (**Fig. 3b, Extended Data Fig. 4a**). We conclude that chromosomes form weak intermingled territories in hESCs, and that as cells differentiate chromosomes become more compact and segregated from each other, consistent with the reduction in inter-chromosomal interactions observed with Hi-C.

We note that at the DE stage, Hi-C data showed that inter-chromosomal interactions remained as high as in hESCs, yet chromosome painting showed that chromosomes were already becoming increasingly territorial. This indicates that relatively high inter-chromosomal interactions as detected by Hi-C do not necessarily imply low chromosomal territoriality. While this could be due to technical limitations, e.g., we may underestimate the size and overlap of the chromosome territories by FISH, it is possible that the process of territory formation involves an intermediary step where chromosomes become less intermingled yet retain extensive interaction surfaces with each other (e.g., at the DE stage), followed by further compaction and increased space between chromosomes at later stages.

To explore inter-chromosomal interactions in another way, we generated aggregated inter-chromosomal Hi-C contact maps (normalized to expected frequencies, see Methods) for all or subsets of chromosomes (**Fig. 3c, Extended Data Fig. 4b**). We observed enriched inter-chromosomal contacts between centromeres (pericentromeric regions in the center of the contact map), as well as enriched inter-chromosomal contacts between telomeres (four corners of the contact map). These enriched contacts were found in hESCs and the four subsequent stages, but they changed in frequency. First, we found that contacts between pericentromeric regions decreased significantly during the ES to DE transition. This was followed by a robust recovery of interaction frequencies from the HB stage onward (**Fig. 3c,d**). Furthermore, we observed that chromosomes exhibit size-dependent differences in this dynamic behavior. Large chromosomes (chr1–10), which on average reside closer to the nuclear periphery in differentiated cells ^46^ generally exhibited lower interchromosomal pericentromeric interaction frequencies than smaller chromosomes (chr11-22) (**Fig. 3c,d; Extended Data Fig. 4b,c**). Large chromosomes displayed relatively weak pericentromeric interactions at distances up to ∼10 Mb (**Extended Data Fig. 4c**), which decreased during the ES to DE transition. For these chromosomes, pericentromeric interactions remained relatively infrequent until the iHLC stage when interactions increased again. In contrast, for smaller chromosomes (chr11–22), pericentromeric interactions were more frequent at all stages, peaking within a narrower genomic neighborhood (∼6 Mb) and remaining elevated up to 12–14 Mb (**Extended Data Fig. 4c**). Pericentromeric interactions between this set of chromosomes decreased at the DE stage and then significantly increased from the HB stage onwards (**Fig. 3d**).

We mapped the subnuclear localization of centromeric regions by co-staining the centromeric protein CENP-B with H3K9me3 (**Fig. 3f**). As expected, bright H3K9me3 puncta co-localized with CENP-B, confirming its enrichment at centromeric and pericentromeric heterochromatin. Centromeric signals were largely scattered throughout the nucleoplasm in hESCs but migrated to the nuclear lamina during the ES to DE transition (**Fig. 3f,g**). This peripheral relocation was transient and unique to the DE stage; by the HB and subsequent HLC stages, a substantial fraction of the centromeric signal had relocated toward the nuclear interior (**Fig 3f,g**). This observation aligns with the inter-chromosomal pericentromeric interactions analysis, where from ES to DE all chromosomes displayed decreased pericentromeric interactions when they reside near the periphery, and gained interactions from the HB stage onward when a number of centromeres relocalize back towards the nuclear interior.

Interchromosomal interactions between telomeric regions displayed different dynamics. (**Fig. 3e, Extended Data Fig. 4d**). In both ES and DE cells chromosomes displayed relatively infrequent telomeric interactions. During the DE-HB transition, contacts increased significantly, decreased somewhat at the iHLC stage, and became more frequent again at the HLC stage. These dynamics were observed for both the set of large and the small chromosomes.

These analyses demonstrate global changes in chromosomes during differentiation, with chromosomes becoming increasingly territorial, centromeres de-clustering and re-clustering while moving towards and away from the periphery, and with telomeres increasingly interacting with each other.

### Compartmentalization strengthening correlates with altered speckle morphology and nuclear lamina composition

The IPT analysis shows that the major increase in compartmentalization during the DE to HB transition is driven mostly by segregation of T.a1, a speckle-dominated sub-compartment, and T.inactive, which includes domains associated with the nuclear periphery. Therefore, we assessed the status of a nuclear speckle component (SRRM2), Lamin B2, and Lamin A/C, and the Lamin B receptor (LBR) during hESC differentiation.

hESCs form very cloud-like, diffuse speckles whereas SRRM2 signal distribution was more uniform throughout the nucleus (**Fig. 4a**). This makes identification of individual speckles through image segmentation in hESCs difficult compared to other stages that have larger speckles with sharper, more distinct boundaries. SRRM2 signal became sharper leading to numerous small speckles after cells transition from hESC to DE and we saw a progressive increase in speckle volume and reduction in number during the HB to HLC transitions, suggesting fusion and consolidation of speckles, leading to structures reminiscent of typical speckles observed in differentiated cells ^47^ (**Fig. 4b,c**). These results show dynamic changes in nuclear speckle morphology during differentiation. This is especially pronounced during the DE to HB transition.

**Fig 4:**
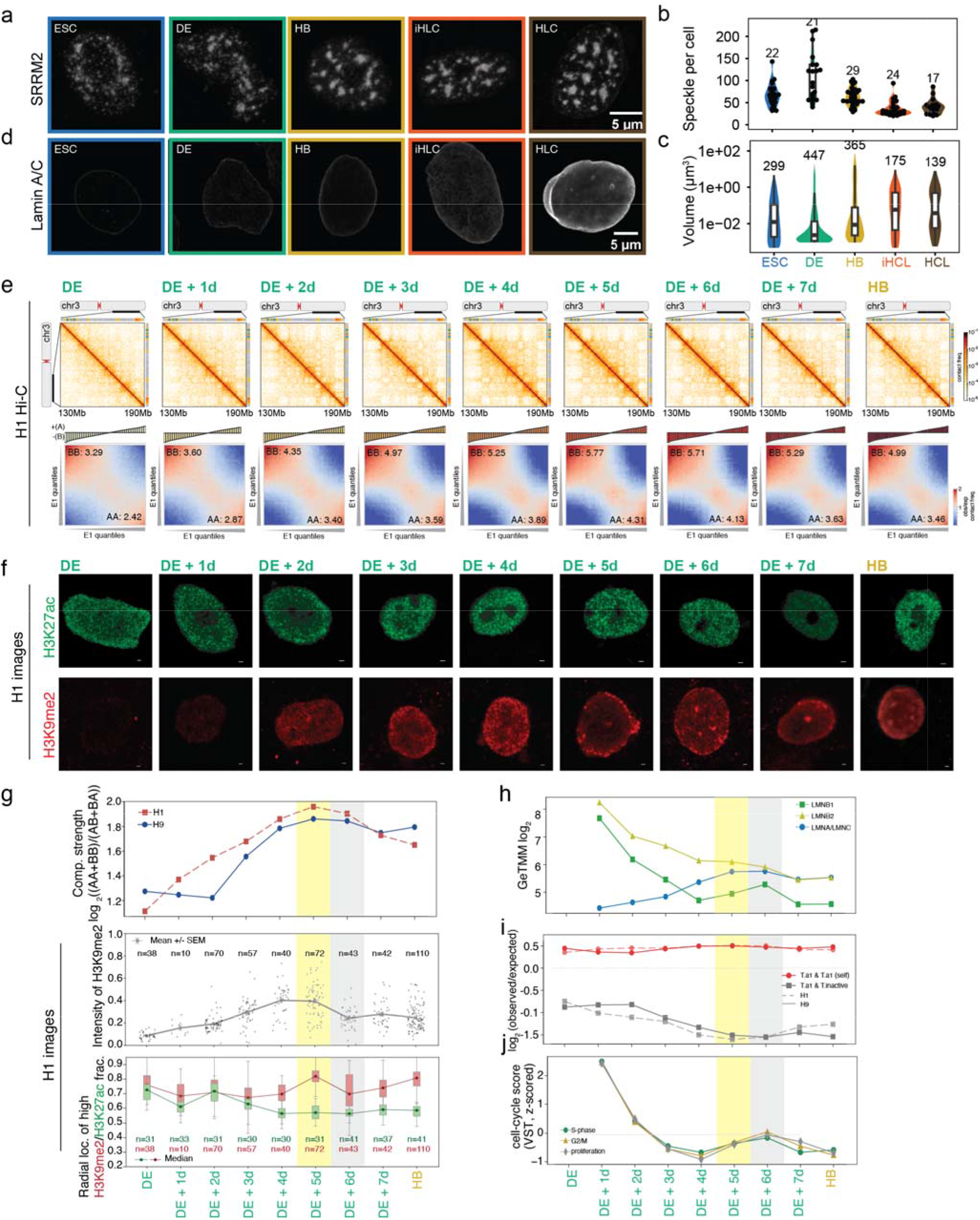
Compartmentalization correlates with changes in speckle morphology and nuclear lamina composition. **a**, 3D projections of cells stained with SRRM2 across 5 stages. Scale bar: 5 μm. **b-c**, Quantifications of speckle volume (b) and Speckle/cell (c) for at least 17 cells for timepoints across hepatic differentiation (ESC = 22, DE = 21, HB = 29, iHLC = 24, HLC = 17). **d**, Representative slices showing lamin A/C staining of cells across 5 stages. Scale bar: 5 μm. **e**, Top: Representative regions (chr3: 130Mb-190Mb) of Hi-C contact frequency maps from 8-day DE-to-HB transition. Bottom: PC1 saddle plots from bulk Hi-C showing compartment strength across 8-day DE-to-HB transition. The strength of AA compartment and BB compartment are labeled on the right bottom and the left top of the saddle plot (top 20% bins are used to calculate the strength). **f**, Representative immunofluorescence images of H3K27ac (green, top row) and second row H3K9me2 (red, bottom row) across 8-day DE-to-HB transition. Scale bar: 10μm. **g**, Quantifying Hi-C and immunofluorescence imaging features. First row: log scaled compartment strength ((AA+BB)/(AB+BA)) across 8-day DE-to-HB transition. Second row: Intensity of H3K9me2 quantified from H3K9me2 immunofluorescence staining across 8-day DE-to-HB transition. Third row: relative radial distance of H3K9me2 (red boxplots) and H3K27ac (green boxplots) high intensity domains in nuclei across 8-day DE-to-HB transition. **h**, Relative gene expression level of lamina components during 8-day DE-to-HB transition in H9 cell lines, including LaminB1, LaminB2, and LaminA/C. **i**, Contact strength quantified from Hi-C discrete-saddle plots of T.a1 self-interactions (red lines) and interactions between T.a1 and T.inactive (gray lines) across the 8-day DE-to-HB transition. Dashed lines indicate H1 cell lines, and solid lines indicate H9 cell lines. **j**, Relative gene expression levels of cell cycle-related genes across the 8-day DE-to-HB transition in H9 cell lines.

Lamin B1, B2, and LBR were expressed mostly in hESCs and DE and then were greatly reduced as detected by RNAseq (**Extended Data Fig. 5a**) and Western blotting (**Extended Data Fig. 5b**). Furthermore, LBR and PRR14 increased from hESC, peaking at DE, which coincides with centromere repositioning at the nuclear periphery. Subsequently, Lamin A/C strongly increased from the HB stage onward and became strongly enriched at the nuclear periphery (**Fig. 4d**, **Extended Data Fig. 5a, b**). These analyses show a correlation between an increase in compartmentalization through segregation of speckle-associated chromatin from peripheral chromatin with cytological changes in speckle morphology, and a change in lamina composition at the nuclear periphery.

### Compartmentalization strengthening correlates temporally with deposition and peripheral localization of H3K9me2

The Hi-C analysis revealed a marked increase in compartmentalization strength during the DE-to-HB transition. Given that this transition occurs over 8 days, we sought to determine changes in compartmentalization at increased temporal resolution by profiling this developmental window at day-by-day resolution using Hi-C and imaging. In both male H1 and female H9 hESC differentiation trajectories, overall compartment strength increased greatly between the DE and HB stages: compartment strength progressively increased from DE through day 5 of the transition (DE+5), peaked there, and subsequently declined before stabilizing at the HB stage (**Fig. 4e,g; Extended Data Fig. 6a,c**). This dynamic pattern in chromosome organization was specific to compartment strength (**Extended Data Fig. 6a-c; Extended Data Fig. 7a-c**). Other features detected by Hi-C including chromatin loop frequencies and TAD insulation scores displayed different temporal changes, further highlighting the complexity of this major differentiation transition.

Our IPT analysis (**Fig. 2**) revealed that both the global accumulation of the heterochromatic mark H3K9me2 (defining the T.inactive trajectory) and the intensification of T.a1 trans interactions occurred most prominently during the DE-to-HB transition (**Fig. 2b, Extended Data Fig. 2b, c**). Therefore, we used immunofluorescence to determine the abundance and subnuclear positioning of these two representative chromatin domain types across the 8-day DE-HB transition. We imaged H3K9me2 along with the active chromatin marker H3K27ac (T.a1–3), at each day of the 8-day transition (**Fig. 4f,g**). We found that H3K9me2 was barely detectable at the DE stage but then became robustly deposited at DE+2 days. Initially, H3K9me2 staining was scattered throughout the nucleoplasm rather than enriched at the nuclear periphery, contrasting with its canonical lamina-associated distribution in pluripotent or mature somatic states ^23^ (**Fig. 4f,g**). In subsequent days the H3K9me2 signal continuously intensified and migrated to the nuclear periphery, reaching maximum peripheral localization at DE+5 (**Fig. 4g**). Active chromatin marked by H3K27ac was present at all days and progressively localized toward the nuclear interior (**Fig. 4f, g, Extended data Fig.5c, d**). These imaging data show that during the 8-day transition, active and inactive chromatin became increasingly spatially segregated, coinciding with the maximum compartmentalization strength detected by Hi-C at DE+5 (**Fig. 4f,g**).

For nuclear speckles, the subnuclear hubs where the highly active T.a1 loci reside, we found a progressive reduction in total counts alongside an increase in individual puncta size during the 8-day transition, reaching a plateau at DE + 5 (**Extended Data Fig. 5e, f**). As shown earlier, there is a transition from expression of lamin B to lamin A/C across the five stages (**Extended Fig. 5a, b**) that exhibits the steepest change at the DE to HB transition. During this transition, the lowest and highest levels of Lamin B and Lamin A/C, respectively, occurred near DE+5d (**Fig. 4h**). In agreement with these metrics, our contact frequency maps showed that trans interactions between T.a1 and T.inactive reached their lowest frequency at DE+5, while self-interactions within the T.a1 trajectory peaked at DE+4 (**Fig. 4i**)—coinciding with changes in nuclear speckle morphology (DE+5) and the maximum spatial segregation of chromatin marked with H3K9me2 and H3K27ac (DE+5). Collectively, these findings suggest that the establishment of genome-wide compartmentalization is driven by the molecular recognition of active versus inactive genomic identities, which subsequently guides their physical sorting into distinct nuclear neighborhoods. Interestingly, genes associated with mitosis and proliferation peak at DE+6 just as compartmentalization strength begins to reverse. This suggests that a higher propensity of cells to divide toward the end of the 8-day transition period modestly counteracts the strengthening checkering patterns in population contact maps (**Fig. 4j**).

Last, we explored the subnuclear localization of centromeric regions across the different chromosome size classes (**Extended Data Fig. 6d, e, f, Extended Data Fig. 7d, e**). Immunofluorescence imaging of H3K9me3-marked centromeric domains revealed that their overall spatial distribution throughout the nucleus largely mirrored the distribution of the H3K9me2 enrichment regions, reaching peak peripheral localization at DE+5 (**Extended Data Fig. 6d, e**). Pericentromeric interactions of large (chr1–10) and small (chr11–22) chromosomes displayed broadly similar, non-monotonic dynamics. Interaction scores initially increased from DE to DE+1, subsequently decreased to levels comparable to or below those at DE, and then increased again around DE+4 before declining modestly toward the HB stage. This pattern was observed for both chromosome size classes, although the increase for small chromosomes did not reach significance in H9 cells (**Extended Data Fig. 6d, f; Extended Data Fig. 7d, e**). Interchromosomal interactions between telomeric regions also followed similar dynamics in large and small chromosomes and more closely paralleled the changes in compartment strength, progressively increasing from DE to DE+5, where they reached a maximum, and subsequently declining toward the HB stage (**Extended Data Fig. 6f; Extended Data Fig. 7e**).

### Chromatin undergoes IPT-specific and genome-wide changes in interaction stability during differentiation

Compartmentalization is thought to be driven by homotypic affinities between chromatin domains sharing a similar epigenetic state. While the increase in compartmentalization observed during differentiation could be driven by association of different domains with different subnuclear structures (speckles, lamina), increased affinities between chromatin domains themselves may also contribute. Chromatin interaction stability, especially chromatin interaction lifetime, can be quantified by liquid chromatin Hi-C ^48^. In this assay, unfixed nuclei are predigested with a restriction enzyme (DpnII) which results in progressive dissolution of chromatin interactions, as chromatin becomes “liquified”. After up to several hours of incubation, cells are crosslinked and Hi-C is performed to determine which chromatin interactions are lost and which ones remain as compared to mock-digested nuclei. Stable, relatively long-lived interactions remain detectable, while unstable, short-lived interactions are lost. This can be quantified by calculating the relative loss of interactions between pairs of loci by the Loss of Structure (LOS) metric ^48,49^ (Methods). LOS quantifies the relative loss of short-range interactions (between loci separated by up to 2 Mb) over the total number of interactions for each genomic bin (150 kb). In this way, a genome-wide profile of chromatin interaction stability is obtained that can be related to compartment status and any other chromatin features. This metric is sensitive to the average fragment size as shorter fragments lead to more dissolution of interactions and higher LOS scores ^48^. Therefore, for each experiment we assess the extent of predigestion by performing fragment size analysis (Agilent Fragment analyzer) and by DpnII-seq ^48^. DpnII-seq data can then be used to remove any such cutting frequency bias in LOS calculations.

LC-Hi-C experiments for each of the hESC, DE, HB and HLC stages were done in duplicate and LOS values generated between replicates were highly correlated (**Extended Data Fig. 8a**). We first applied LC-Hi-C to hESC nuclei (**Extended Data Fig. 8b**). Using the standard protocol we developed before for differentiated cells ^48,49^, we noticed that chromatin in hESCs dissolved faster than in K562, U2OS (**Extended Data Fig. 8c**) and HepG2 cells (**Extended Data Fig. 8d**). Further, each stage of the hepatic differentiation generated different fragment size distributions where fragments become progressively larger throughout differentiation (**Extended Data Fig. 8e**). The faster dissolution of chromatin interactions in hESCs could be the result of the more efficient digestion leading to shorter chromatin fragments whose interactions are expected to be lost faster (higher LOS). In addition, the rapid loss of interactions could be due to generally unstable interactions in these cells. Therefore, to enable direct comparison of chromatin interaction stability in hESCs and differentiated cells, we performed paired experiments where hESC nuclei and nuclei from DE, HB or HLC were subjected to LC-Hi-C. We varied the amount of DpnII for pre-digestion of nuclei followed by a one-hour incubation that produced very similar chromatin fragment length distributions as assessed by Agilent Fragment analyzer analysis (**Extended Data Fig. 8f,g**). hESCs nuclei required lower concentrations of DpnII, to generate similar fragment length distributions, indicating hESC chromatin is more open and readily digestible, as reported before (ES cell chromatin state reviewed in ^16^). We then plotted observed/expected LC-Hi-C contact maps (**Fig. 5a**), and calculated LOS values genome-wide for each pairwise comparison (**Fig. 5b**). We determined LOS value distributions per IPT (**Fig. 5c**). Finally, we corrected LOS for any regional variation in digestion efficiency using matched DpnII data (**Fig. 5d**; LOS residuals, see ^48^ and Methods).

**Fig. 5:**
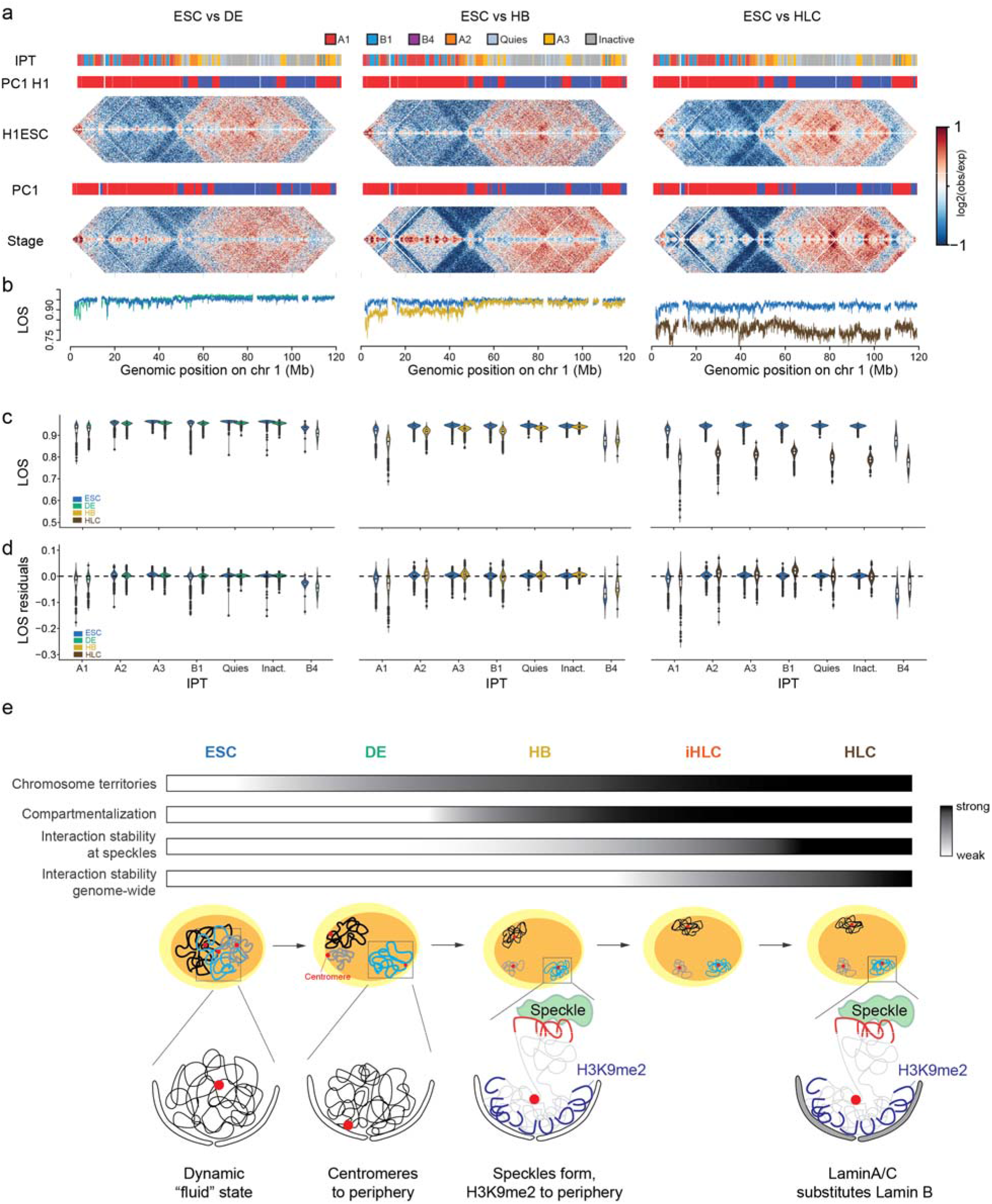
Sequential stabilization of speckle-associated and global chromatin interactions. **a**, Log2(observed/expected) contact frequency heatmaps (binned at 250kb) aligned with tracks of the chromosome 1 p-arm showing SPIN states (first bar), PC1 (second bar), and LOS (**b**) for ESC/DE (left), ESC/HB (middle) and ESC/HLC (right) fragment size distribution-matched pairs. Violin plots displaying LOS (**c**) and LOS residuals (**d**) by IPT for the same pairs are shown. **e**, Model depicting the changes in chromatin organization and chromatin interaction stability across in vitro hepatic differentiation. LOS data is binned at 150kb.

Several insights were obtained with these analyses. First, we found that LOS was higher in hESCs than in HB and HLC. This indicates that chromatin interactions dissolve faster in hESC than in these differentiated cells, even though chromatin was digested to similar length distributions (**Fig. 5b,c**, second and third column; **Extended Data Fig. 8g**). This was observed genome-wide and for all IPTs indicating that chromatin interactions are generally more unstable in hESCs (**Fig. 5c**). Second, LOS in DE was similar to that for hESCs (**Fig. 5b-c**; first column). This indicates that chromatin interactions are as unstable in DE as they are in hESCs. Third, chromatin interactions became more stable and longer-lived at the HB stage and this was most pronounced for T.a1 speckle associated chromatin. Interactions within T.quies and T.inactive domains stabilized only at the HLC stage when interactions throughout the genome became much more stable, including for IPTs not associated with speckles or lamina (e.g. T.active 2 and 3). **Fig. 5b** illustrates these changes in chromatin interaction stability for a 120 Mb region along chromosome 1. LOS along this region was high in both hESC and DE, while the LOS decrease in HB was specific to a large domain that contains mostly T.a1 chromatin. In a flanking region, containing mostly T.inactive chromatin (second column), LOS remained as high as in hESCs. Finally, LOS decreased along the entire region in HLCs compared to a matched hESC LOS profile (third column). This is evident in the contact frequency maps wherein interactions are largely preserved along the diagonal after DpnII digestion in HLC compared to its paired ESC sample (**Extended Data Fig. 8h**). This preservation of contacts is also evident when plotting average contact frequency by distance (**Extended Data Fig. 8i**).

Analysis of LOS residuals allows determining relative stability of interactions in different IPTs within each cell stage (**Fig. 5c**). This analysis showed that T.a1 and T.b4 heterochromatin were more stable relative to other IPTs in the same cell type. We described similar results in a recent study where we find that in K562 cells the longest-lived chromatin interactions in the genome occur in speckle-associated and lamin-associated domains ^49^. These results show that chromatin in hESCs and DE is quantitatively less stable, shorter lived and therefore biophysically distinct from differentiated cells. This may contribute to the weaker compartmentalization observed above. Interestingly, unstable interactions are not a feature unique to hESCs, as DE cells show similar unstable contacts, and similarly weak compartmentalization. As cells differentiate to HB, chromatin interactions start to stabilize genome-wide but this occurs first at T.a1, followed by genome-wide stabilization at the HLC stage. These dynamics correlate with increased compartmentalization strength, formation of large nuclear speckles, and an exchange of Lamin B1/2 with Lamin A/C at the periphery.

## Discussion

We describe a multi-step pathway by which nuclear organization converts from a pluripotent to a differentiated state (**Fig. 5e**). First, chromosomes unmingle, correlated with decreased histone acetylation, while remaining weakly compartmentalized. Then, as they commit to the hepatocyte lineage, chromosomes increasingly separate and compartmentalize in a manner that correlates with increased H3K9me2 deposition, differentiation of nuclear speckle morphology, and alteration in lamina composition. Strikingly, at this stage chromatin interactions become more stable, especially at speckle-associated domains. Finally, during maturation, chromatin interactions become more stable genome wide.

In contrast to most other cell types, we find that in hESCs, chromosomes appear to lack territoriality. By chromosome painting, they appear partitioned into separate domains that intermingle with each other extensively. Similar observations were reported by Gelali and co-workers, who compared territoriality in human ESCs to that in human IMR90 cells. They found that chromosome territories in hESCs intermingle much more frequently and to a larger extent than in IMR90 cells ^18^. It is not known how chromosomes form separate territories in differentiated cells. It has been proposed that, in dividing cells, there is insufficient time for chromosomes to mix after mitosis ^50^.

Surprisingly, we find that chromosomes are most intermingled in hESCs – which have the shortest cell cycle – and as cells differentiate and the cell cycle slows, territoriality increases. This implies that after chromosomes segregate during mitosis in hESCs, they rapidly and globally decondense upon mitotic exit. Indeed, hESC interphase chromosomes become so diffuse that greater concentrations of chromosome paint probes are required to detect them via FISH (see Methods). We further note that relative to DE and further differentiated cells, the more acetylated chromatin in hESCs may contribute to their decondensed state (**Extended Data Fig. 5e,f**). Consistently, hyperacetylation from histone deacetylase inhibitors, leads to more extensive mixing between chromosomes as observed using microscopy ^51^ and increased interchromosomal interaction as detected by Micro-C ^52^. In differentiated cells, chromatin decondensation after mitotic exit must be either less extensive or other processes must act to prevent intermingling and keep chromosomes individualized in interphase.

By Hi-C we observed a higher level of interchromosomal interactions in hESCs than in differentiated cells (i.e., from the HB stage onwards), while interchromosomal interactions in DE cells were comparable to those in hESCs. Intriguingly, chromosomes already began to appear more compacted and less admixed during the hESC-to-DE transition, i.e., they become more territorial at an earlier stage than global *cis* vs *trans* contacts would naively suggest. This implies that while chromosomes appear more individualized at the DE stage, they maintain a large surface area of interaction. This could be related to the fact that during the hESC-to-DE transition nuclear morphology changes from a round to a more flattened shape. In flattened nuclei, chromosomes may be well separated yet lie as disks on top of each other, preserving relatively large areas of contact. As DE cells transition to the HB stage, interchromosomal interactions decrease and remain similarly low during subsequent stages, while territoriality continues to increase. This again indicates that the level of interchromosomal interactions is not a quantitative measure of territoriality. Nuclear bodies, such as nuclear speckles, and lamina-associated heterochromatin, both of which become prominent and well-defined during the DE to HB transition, may contribute to the increased territoriality seen by imaging and the reduced inter-chromosomal contacts seen by Hi-C.

Our data show that the development of strong chromosome territoriality involves multiple steps where chromosomes that are initially extensively intermingled in hESCs first unmingle at the hESC-to-DE transition but maintain frequent interchromosomal interactions before becoming increasingly separated as cells further differentiate. While chromosome territoriality starts to increase, centromeres move to the nuclear periphery. Whether this contributes to territory formation is not known. Clearly, how chromosome territories form, and how cells maintain this state remains a major question in the field of nuclear organization. Our study shows that hESC differentiation provides an experimental system to explore this long-standing question.

Compartmentalization is weak in hESCs and DE cells and increases as DE cells transition to HB cells. This is detected by Hi-C, reflected in a weak checkerboard pattern, and by microscopy as a co-localization of domains marked by active or inactive chromatin modifications (i.e., H3K27ac and H3K9me2). Combined, this strongly implies that weak compartmentalization detected by population-averaged Hi-C data is not due to large cell-to-cell variation in genomic positions of active and inactive chromatin domains in hESCs and DE cells. If that were the case, microscopy would still detect segregation of H3K27ac and H3K9me2 marks at the single cell level.

Compartmentalization is thought to be driven by affinities between chromatin domains of similar type, leading to phase separation ^4,48,53^. This indicates that homotypic affinities increase during differentiation, possibly due to changes in chromatin composition and histone modifications. Interestingly, the increased compartmentalization strength detected by Hi-C and by immunofluorescence during the DE-to-HB transition coincides precisely with a strong increase in deposition of the H3K9me2 mark. Many other modifications may, of course, contribute as well.

Rather than assigning each locus a compartmental identity at each stage and matching assignments post hoc, we decomposed *trans* interaction profiles from all five stages into a single joint basis and aligned the score vectors so that each 50 kb bin is described by a single spectral trajectory. Clustering and characterizing these trajectories yielded seven IPTs identifying broad regions of the genome that remain relatively architecturally stable during differentiation (T.a1, T.a2, T.a3, T.b4, T.quies) as well as other broad regions that are significantly remodeled, architecturally and epigenetically, in a concerted manner over the five stages due to heterochromatin accumulation (T.pcg, T.inactive). This approach is also sensitive to features that are spectrally invisible at any single stage. For example, Polycomb-associated chromatin is too narrow and sparse to emerge as its own compartment in hESCs, yet its constituent bins change coherently over differentiation and are recovered as part of a distinct IPT.

Increased compartmentalization strength also coincides with formation of large nuclear speckles, and a change from Lamin B to Lamin A/C at the nuclear periphery. Speckles associate with highly active chromatin, and inactive chromatin associates with the nuclear lamina. It is possible that the differentiation of speckles and the change in lamina composition contributes to better spatial segregation of active and inactive chromosomal domains that associate with these structures. While appealing, several lines of evidence suggest this is too simplistic. Previous work has shown that loss of anchoring of heterochromatin from the periphery, e.g., in inverted nuclei of mouse rod cells, does not lead to loss of compartmentalization and that interactions between heterochromatic domains do not depend on lamina association ^53,54^. Also, we recently showed that formation of stable long-lived chromatin interactions between highly active loci that are normally associated with nuclear speckles, do not depend on association of these loci with the nuclear speckle itself ^49^. Therefore, we propose that compartmentalization as detected by Hi-C is driven mostly by chromatin intrinsic factors, and the association of chromatin compartments with sub-nuclear structures such as speckles and the periphery is a secondary, and possibly mechanistically distinct phenomenon.

Finally, chromosome conformation in hESCs is distinct from that in differentiated cells by being driven by short-lived chromatin interactions. Others have shown that chromatin in pluripotent cells is much more mobile than in differentiated cells ^15,17^. Possibly this high mobility is facilitated by the fact that interactions between loci are very transient, as detected by LC-Hi-C. This low chromatin interaction stability and high mobility may lead to mixing of chromatin domains and contribute to low territoriality and weak compartmentalization. As cells differentiate chromatin interactions become more long-lived globally. Interestingly, using an entirely different approach Zidovska and co-worked used imaging to track chromatin mobility in mouse embryonic stem cells, and its differentiated progeny and found that chromatin in stem cells chromatin behaves like a Maxwell fluid ^55^. As cells differentiate, chromatin was found to undergo a phase transition where chromatin became less mobile and formed a mix of fluid-like and solid-like phases. The biophysical basis of this phenomenon is currently not known but likely again involves general changes in chromatin composition.

At all cell stages, loci associated with nuclear speckles (T.a1) and heterochromatic domains (T.b4) are engaged in relatively more stable chromatin interactions. At the DE-to-HB transition, chromatin interactions start to stabilize genome-wide, but interactions at speckle associated domains stabilize the most. Heterochromatic interactions stabilize only at the HLC stage, along with global stabilization. This suggests that interactions along and between speckle-associated domains are stabilized through distinct mechanisms as compared to the rest of the genome. Consistent with this model, we recently showed that chromatin interactions at speckle-associated domains are RNA-dependent, while chromatin interaction stability in throughout most of the rest of the genome (except nucleolus-associated domains) is not ^49^.

Here we focused on compartmentalization, chromosome territoriality and nuclear organization. At the level of chromatin looping, as detected by HiCCUPS loop calling ^56,57^, we did not detect major differences between pluripotent cells and differentiated cells: loops increase in number, with many loops shared among the 5 cell stages as reported for other cell differentiation systems (e.g., ^57–60^), and loops that did change were correlated with cell stage-dependent gene expression changes, compartment changes and RAD21 occupancy. Thus, it appears that at the level of chromatin loops, pluripotent cells and differentiated cells are not distinct in ways other than expected for any two different cell types.

The mechanisms by which cells establish and maintain chromosome territories and spatially compartmentalize different types of chromatin domains remain surprisingly poorly understood even though these phenomena have been studied for decades. The hESC differentiation pathway we characterize here in detail at the genomic and cell biological level provides a genetically tractable system where these features of nuclear organization undergo dynamic changes at distinct differentiation transitions. This provides an experimental opportunity to start to dissect the molecular and biophysical mechanisms driving these major events.

## Methods

### Cell culture

#### Human Embryonic Stem Cell (hESC) Culture

Human embryonic stem cells (hESCs; H1 line, WiCell, WA01, lot no. WB35186; H9 cell line, WiCell, WA09) were maintained under feeder-free conditions on plates coated with Matrigel hESC-Qualified Matrix (Corning, 354277). Cells were cultured at 37°C in a humidified incubator with 5% CO2 and fed daily with fresh mTeSR1 medium (StemCell Technologies, 85850). Routine passaging was performed every 4–5 days using ReLeSR reagent (StemCell Technologies, 05872). For single-cell dissociation prior to downstream assays or differentiation, cells were treated with TrypLE Express (Thermo Fisher, 12604013).

#### Differentiation of H1 Cells to Hepatocyte-Like Cells (HLCs)

Directed differentiation was conducted by adapting and optimizing an established hepatic differentiation protocol for human pluripotent stem cells ^61^ for compatibility with the protocol generating the first two stages (hESC, definitive endoderm) which was developed as part of 4DN Phase 1 ^36^, and to provide more homogeneous differentiation material.

Briefly, H1 and H9 hESCs were dissociated into a single-cell suspension using TrypLE Express and seeded onto growth factor-reduced (GF−) Matrigel-coated plates (0.125 mg/ml) at a density of 1×10^6^ cells/ml. **Definitive Endoderm (DE) Induction:** DE differentiation was carried out over 4 days using the STEMdiff Definitive Endoderm Kit (StemCell Technologies, cat#5110). On day 2, cells were washed with DMEM/F12 (ThermoFisher cat#11320-033), and the medium was replaced with STEMdiff DE Basal Medium supplemented with Supplement with MR+CJ. On days 3 and 4, cells were fed daily with basal medium containing Supplement CJ only. Successful DE induction was validated via immunofluorescence staining for SOX17 (R&D Systems, AF1924) and FOXA2 (EMD Millipore, 07-633). **Hepatic Specification (Hepatoblasts):** To initiate hepatic specification, DE cells were detached using TrypLE and re-seeded at a density of 1.5×10^6^ cells/ml onto GF(−) Matrigel-coated plates. Cells were cultured in a hepatic differentiation basal medium [High-glucose DMEM/F12, 10% KnockOut Serum Replacement (KOSR, ThermoFisher cat#10828-028), 1% L-glutamine, 1% non-essential amino acids (NEAA, ThermoFisher cat#11140-050), and 1% Penicillin/Streptomycin supplemented with 100 ng/ml hepatocyte growth factor (HGF; Peprotech cat# 100-39H), 1% DMSO (Sigma-Aldrich, cat#D2650), and 10 μM Rho-associated protein kinase (ROCK, Stemgent cat#04-0012-010) inhibitor Y-27632. Twenty-four hours post-seeding, the medium was replaced with differentiation basal medium containing 100 ng/mL HGF (without ROCK inhibitor) and refreshed daily for 7 days. The resulting hepatoblast (HB) population was characterized by expression of TBX3 (Santa Cruz Biotechnology, sc-166623) and AFP (Santa Cruz Biotechnology, sc-8399). **Immature hepatocyte-like cells:** To drive hepatoblasts toward immature hepatocyte-like cells, the medium was transitioned to differentiation basal medium supplemented with 10^−7^ M dexamethasone (Sigma cat#D4902, replacing HGF). The medium was changed daily for 3 days. Differentiation was confirmed by positive staining for albumin (ALB; Sigma, A6684) and AFP. **Hepatocyte-like cells:** For final maturation into HLCs, the medium recipe are: William’s E Media, 10% FBS, 1% Penicillin/Streptomycin, 1.8% DMSO, 1ug/ml insulin (Sigma cat#I92780, supplement with 4.8ug/ml hydrocortisone (Sigma cat#H4001). Cells were maintained in this maturation medium for an additional 12 days with media changes every 2-3 days. Mature HLCs were characterized by the expression of HNF4a (Abcam, ab41898) and ALB.

### Hi-C Library preparation

Hi-C libraries were generated following the 4DN Hi-C 3.0 protocol ^62,63^ with minor modifications. Briefly, undifferentiated and differentiated cells were harvested at a density of approximately 5×10^6^ cells per plate, washed twice with 10 ml of HBSS, and fixed according to the protocol guidelines. Cell pellets collected from the various differentiation stages were snap-frozen in liquid nitrogen and stored at -80°C. Upon collecting all time points, the crosslinked cell pellets were thawed, resuspended in Hi-C lysis buffer, and subjected to overnight genomic digestion at 37°C using a dual-enzyme cocktail of DpnII and DdeI (NEB). The resulting overhanging DNA ends were filled in with biotin-14-dATP (Life Technologies) at 23°C for four hours, followed by proximity ligation using T4 DNA Ligase (Life Technologies) at 16°C for four hours. To ensure complete reversal of crosslinks, Proteinase K (Thermo Fisher) was added twice to the samples during an overnight incubation at 65°C. Purified DNA ligation products were sonicated to a size range of 200–350 bp, and biotin-tagged fragments were captured using Dynabeads MyOne Streptavidin C1 (Invitrogen). A-tailing and Illumina adapter ligation were performed directly on the bead-bound DNA, and the libraries were amplified using NEBNext Multiplex Oligos for Illumina (Dual Index Primers, NEB). After removing primer dimer contamination using a 1.1× AMpure XP bead cleanup, the final libraries were sequenced on an Illumina NovaSeq 6000 platform using a paired-end 50 bp (PE50) read recipe.

### ChIP-seq

Briefly, for each stage, 2-15 x10^6^ of cells were crosslinked with 1% formaldehyde. The cell pellet was resuspended in lysis buffer (20mM TrisHCl pH8.0, 85mM KCl, 0.5% NP-40 + protease inhibitor), incubate 15min on ice, centrifuge 5min 4°C, resuspend pellet in sonication buffer (20mM Tri-HCl pH 8.0, 0.2% SDS, 0.5% sodium deoxycholate, and protease inhibitor). The chromosomes were sonicated to fragments around 200–500bp using BioRaptor Pico (30sec ON, 30sec OFF, 7-8 cycles). After spinning at 16,000g for 10min at 4°C, the fragmented chromatin in the supernatant was split into 4 aliquots per condition and diluted in 1200ul IP buffer with final 0.1% SDS concentration (20mM Tri-HCl pH 8.0, 150mM NaCl, 2mM EDTA, 0.1% SDS, 1% Triton-100, and protease inhibitor). For each tube, around 1-5ug of antibody was added. The primary antibodies used in this study included, CTCF (Cell Signaling, #3418S), RAD21 (Abcam, #ab992, recognizes C-terminal of RAD21), RAD21 (Abcam, #ab154769, recognizes N-terminal of RAD21), and Rabbit IgG (Sigma, #I-5006). Chromatin was incubated with primary antibodies on a rocker at 4°C for overnight. After rewashing with IP buffer, 20ul of Dynabeads Protein A (Thermo Fisher, #10002D) were added to each tube followed by incubation on a rocker at 4°C for 4 hours. After the beads were washed for three times, immunoprecipitated DNA was eluted and 5-10ng ChIP DNA was used to prepare sequencing libraries in the same way as described for Hi-C above.

### Cut&Run

Cells were washed once with ice cold DPBS and dissociated into single cells using TrypLE Express. 500,000 cells were resuspended fresh in ice cold PBS for each reaction, except for replicate two of HB, iHLC, HLC, which used 100,000 cells per reaction. The library was prepared using CUTANA ChIC/CUT&RUN Kit v3 (Epicypher, 14-1048) and CUTANA CUT&RUN Library Prep Kit (Epicypher, 14-1001) according to manufacturer’s protocol and as previously published ^64^. Digitonin was used at 0.01% final concentration, except for replicate two of HB, iHLC, and HLC, which used at 0.05% final concentration, for cell permeabilization, and 0.5 ug primary antibodies were used for each reaction. Library concentration was quantified using Qubit 1X dsDNA HS Assay Kit (Invitrogen, Q33231). Library quality and fragment size distribution were assessed using Agilent Bioanalyzer with High Sensitivity DNA Kit (Agilent Technologies, 5067-4626).

### ATAC-seq

ATAC-seq was performed on iPSC-derived hepatocytes using the ATAC-seq kit (ActiveMotif, 53150) according to manufacturer’s protocol and as previously published ^65^. Briefly, cells were dissociated using TrypLE Express (Gibco, 12604) and neutralized in culture media. Cells were pelleted at 500 g for 5 min and counted using an automated cell counter. 100,000 cells were subsequently transferred to a new Eppendorf tube and washed once with ice-cold 1X PBS. Cell pellet was then gently resuspended in 100ul ice-cold ATAC-seq Lysis Buffer and transferred to a new PCR tube on ice. The PCR tube was placed in a tube holder and immediately centrifuged at 500 g for 10 min at 4C. After the spin, the supernatant was removed carefully, and the pellet was used immediately for tagmentation reaction and purification. Library concentration was quantified using Qubit 1X dsDNA HS Assay Kit (Invitrogen, Q33231). Library quality and fragment size distribution were assessed using Agilent Bioanalyzer with High Sensitivity DNA Kit (Agilent Technologies, 5067-4626).

### RNA-seq

Stem Cells and differentiated cells were grown into 6-well plates before collection of one well into 1 ml of Trizol and storage at -80°C. Total RNA was isolated from the aqueous phase of TRIzol®. Briefly, per 1 ml of TRIzol®, 0.2 ml of Chloroform Reagent was added and 1.5 ml tubes were vortexed for 30 seconds and centrifuged at 4000 rpm for 5 minutes at 4°C. The aqueous phase was precipitated by adding 0.5 ml of isopropyl alcohol (2-propanol) per 1 ml of TRIzol®. Samples were incubated at room temperature for 10 minutes before centrifugation (14000 rpm for 10 minutes at 4°C). Supernatant was poured off and the RNA pellet washed once with nuclease free 75% ethanol and centrifuged for 5 min at 14000 rpm at 4°C. Precipitated RNA was dissolved in 50 µl of nuclease free water. For the 5 stages of differentiation, RNA was submitted to BGI where it was enriched for mRNA before fractionation and cDNA generation (BGI). The cDNA was subjected to 100 nt paired-end strand-specific mRNA sequencing by DNBSEQ (BGI). For the 8 days of differentiation between DE and HB, we performed on-column DNase Digestion with DNAse I using the Qiagen RNeasy kit (Qiagen). Cleaned RNA was run on a 1% TBE agarose gel and RIN >7 was confirmed by18/28S. We used the NEBNext® rRNA Depletion Kit v2 (E7400L) to remove rRNA and the NEBNext® Ultra™ II Directional RNA Library Prep Kit for Illumina (E7760S) to generate cDNA libraries for 150 bp paired-end (300 cycles) Illumina sequencing on a NextSeq2000 instrument.

### Liquid Chromatin Hi-C

#### Nuclei Isolation

The experimental protocols described herein were adapted from ^48^ with specific modifications. Approximately 50–100 x10^6^ cells (H1ESC, DE, HB, or HCL) were harvested, washed twice with Hanks’ Balanced Salt Solution (HBSS; Thermo Fisher Scientific, Cat# 14025134), and pelleted by centrifugation at 4°C for 10 min. Pelleted cells were resuspended in 15 ml of ice-cold NB buffer [10 mM PIPES pH 7.4, 10 mM KCl, 2 mM MgCl2, 1 mM DTT, 1× Protease Inhibitor Cocktail (Thermo Fisher Scientific, Cat# 78429), and 0.1% NP-40], transferred to a 15 ml Dounce tissue grinder (Kimble, Cat# 885303-0015), and incubated on ice for 10 min. Cell lysis was performed using 20 strokes with an A-type pestle, followed by a 20 min ice incubation and an additional 20 strokes. A 10 μl aliquot of the lysate was mixed with an equal volume of Trypan Blue to evaluate nuclear integrity and complete membrane disruption; if incomplete, an additional 10 strokes were applied.

The 15 ml cell lysate was divided equally and layered onto three sucrose cushion tubes containing 10% sucrose prepared in NB buffer, then centrifuged at 3,500×g for 30 min at 4°C. Supernatants were carefully aspirated to a residual volume of approximately 5 ml. The remaining fractions containing the nuclear pellets were pooled, transferred to a clean tube, and spun at 1,500×g for 5 min at 4°C. After removing the supernatant, the nuclear pellet was resuspended in 2 ml of NSB buffer [50% glycerol, 250 mM sucrose, 1 mM DTT, and 1× Protease Inhibitor Cocktail]. Intact nuclei were quantified using Trypan Blue staining, adjusted to a final concentration of 5×10^6^ nuclei/ml, and stored in 500 μl aliquots at −80°C.

#### Nuclei Digestion and Processing

Hi-C and Liquid Chromatin Hi-C (LC-Hi-C) procedures were adapted from Belaghzal et al. 2017 ^66^ and Belaghzal et al. 2021 ^48^, respectively. Frozen nuclear aliquots were thawed on ice, diluted with 1 ml of HBSS, mixed gently by inversion, and pelleted at 1,500×g for 8 min at 4°C. Nuclei were washed once in 1× NEBuffer r3.1 (NEB, Cat# B6003S), quantified, and resuspended at 6 millions nuclei/ml. DpnII (50 U/μl; NEB, Cat# R0543M) was added to a final working concentration of 0.25 U/μl or 0.5 U/μl for digested conditions, while an equivalent volume of 1× NEBuffer r3.1 was added to control mock-digested samples. Aliquots of 500 μl were distributed into 1.5 ml tubes and incubated at 37°C for 2–4 h. Following incubation, digest aliquots were partitioned for downstream assays: 1. **Fragment Analysis:** 1x10^6^nuclei from both mock- and DpnII-treated samples. 2. **DpnII-seq:** 1x10^6^ nuclei from the DpnII-treated sample only. 3. **LC-Hi-C:** 3x10^6^ nuclei from both mock- and DpnII-treated samples.

LC-Hi-C samples were immediately crosslinked with 1% formaldehyde, washed, flash-frozen, and stored at −80°C. Genomic DNA for fragment analysis was recovered via either standard phenol:chloroform extraction or AMPure XP bead purification.

#### Hi-C procedure

Hi-C libraries were generated following established methods ^66^ with minor modifications. Flash-frozen, crosslinked mock- and DpnII-pre-digested nuclei were subjected to restriction digestion using DpnII (400 U per sample) overnight at 37°C. The resulting sticky ends were filled in with biotin-14-dATP at 23°C for 4 h, followed by blunt-end proximity ligation using T4 DNA ligase at 16°C for 4 h. Protein crosslinks were reversed by overnight Proteinase K treatment at 65°C. Re-ligated DNA was purified, fragmented to an average size of 200 bp via ultrasonic shearing, and size-selected (100–350 bp). Purified fragments underwent end-repair and dA-tailing prior to streptavidin-mediated pull-down of biotinylated junctions. Following Illumina TruSeq adapter ligation, libraries were PCR-amplified, purified to eliminate primer-dimers, and subjected to paired-end sequencing on Illumina platforms (HiSeq 4000, NextSeq 2000, or NovaSeq).

#### DpnII-seq

Isolated DpnII-digested nuclei (1x10^6^) were transferred to a 1.7 ml microcentrifuge tube containing 10 μl of 0.5M EDTA, incubated at 65°C for 20 min, supplemented with 50 μl of 10 mg/ml Proteinase K, and digested at 65°C for 3 hrs to overnight. Genomic DNA was isolated using phenol:chloroform extraction with ethanol precipitation or AMPure XP bead cleanup, followed by elution in 85 μl of TLE buffer.

To end-fill restriction overhangs, 15 μl of a fill-in master mix—comprising 0.5 μl nuclease-free water, 1.5 μl 10×NEBuffer 3.1 (NEB, Cat# B7203S), 0.4 μl each of 10 mM dCTP, dGTP, and dTTP (Invitrogen, Cat# 56173, 56174, 56175), 9.4 μl of 0.4 mM biotin-14-dATP (Thermo Fisher Scientific, Cat# 19524016), and 2.5 μl Klenow DNA Polymerase (5 U/μl; NEB, Cat# M0210L)—was added to each sample. Reactions were incubated at 23°C for 1.5 h, treated with 1 μl of 10 mg/ml RNase A at 37°C for ≥30 min, and diluted to a final volume of 132 μl with nuclease-free water.

After agarose gel quality control, samples were sonicated, size-selected, and end-repaired as described in the LC-Hi-C workflow. Following end-repair, DNA was purified using a 2× AMPure XP bead-to-sample ratio (omitting the streptavidin pull-down at this stage) and eluted in 41 μl of TLE buffer. Samples were dA-tailed, subjected to a 2× AMPure cleanup, and eluted in 32 μl of nuclease-free water. Illumina adapters were ligated by adding 8 μl of 5× Ligation Buffer (Invitrogen, Cat# Y90001) and 10 μl of ligation master mix. Ligated products were purified with 2× AMPure beads and eluted in 90 μl of nuclease-free water.

Each library was supplemented with 10 μl of 10× NEB CutSmart Buffer and 3 μl of USER Enzyme (NEB), then incubated at 37°C for 1 h. Biotinylated fragments were captured using streptavidin magnetic beads as described in the LC-Hi-C protocol, with final wash steps carried out in 1× NEBuffer r2.1 instead of TLE. PCR cycle optimization and library amplification proceeded according to the standard LC-Hi-C workflow, with the ClaI restriction digest step omitted.

### Western blotting

For each stage, approximately 2x10^6^ cells were harvested, washed twice with DPBS, and incubated with 50 μl of RIPA buffer (#89900) containing protease inhibitors and TurboNuclease (#N0103M) for 30 min at 4°C. Lysates were scraped, cleared by centrifugation at 8,000g for 10 min, and collected. Samples were mixed with 4× Laemmli sample buffer (Bio-Rad, #1610747), denatured at 95°C for 5 min, and loaded at cell-equivalent volumes. For anti-Lamin B Receptor blots, samples were instead incubated at 70°C for 10 min to prevent irreversible hydrophobic aggregation. Proteins were resolved using NuPAGE 4–12% Bis-Tris gels (#NP0322BOX) in NuPAGE MES SDS running buffer (#NP0002) and transferred to nitrocellulose membranes (#1620112) at 30 V for 2 h at 4°C in 1× transfer buffer (#35040). Membranes were blocked with 5% milk in TBST for 60 min at room temperature, incubated overnight at 4°C with primary antibodies (1:1,000), and washed three with TBST. Following 2 h incubation with HRP-linked secondary antibodies (1:5,000) at room temperature and three subsequent washes, blots were developed using SuperSignal West Dura substrate (#34076) and imaged on a Bio-Rad ChemiDoc system. Primary antibodies utilized included anti-Lamin A/C (Abcam, ab108595), anti-Lamin B2 (Abcam, ab151735), anti-β-actin (Cell Signaling, 3700S), and anti-Lamin B Receptor (LBR; Thermo Fisher, PA5-66827).

### Imaging

#### Immunofluorescence

IF staining was performed as previously described ^67^ with minor modifications. Briefly, cells cultured on sterile coverslips (ThorLabs, CG15NH) were fixed with 4% paraformaldehyde for 10 min, permeabilized with 0.2% Triton X-100 for 10 min, and blocked with 2% BSA for 30 min at room temperature. Primary antibodies were incubated overnight at 4°C, followed by secondary antibody incubation for 60 min at room temperature; 1× PBS washes were performed between all steps. Nuclei were stained with 1 μM DAPI for 10 min. Coverslips were mounted using ProLong Diamond Antifade Mountant with or without DAPI (Invitrogen, P36962/P36961) and cured for 24 h in the dark.

Primary antibodies included: SOX17 (R&D Systems, AF1924), FOXA2 (EMD Millipore, 07-633), TBX3 (Santa Cruz, sc-166623), ALB (Sigma, A6684), AFP (Santa Cruz, sc-8399), HNP4a (Abcam, ab41898), SOX2 (Santa Cruz, sc-365823), and OCT4 (Santa Cruz, sc-5279), CENPB (Thermo Fisher, PA5-65869), H3K27ac (Cell signaling, CST8173T), H3K9me2 (Abcam, ab1220), H3K9me3 (Abcam, ab8898), LAMINAC (Abcam, ab224816), H1.2 (Abcam, ab4086), SRRM2 (Thermo Fisher, PA5-66827). Secondary antibodies (1:1,000) were goat anti-mouse Alexa Fluor 568 (Abcam ab175473) and goat anti-rabbit Alexa Fluor 488 (Abcam ab15007).

#### Chromosome painting

Cells cultured on coverslips were fixed with 4% paraformaldehyde (PFA) for 10 min at room temperature, washed once with 1× DPBS, and either stored at 4°C or processed immediately. Next, cells were permeabilized with 0.2% Triton X-100 for 10 min, followed by three of 3 min washes in 1× DPBS. Coverslips were then treated with 1 μg/μl RNase A in 1× DPBS for 60 min at 37°C to remove cellular RNA, rinsed with nuclease-free water for 1 min, and dehydrated through a graded ethanol series (75%, 85%, and 100% for 3 min each). Following complete air-drying, a 15 μl probe mixture containing 30 or 50 pM whole-chromosome painting probes (H1-hESC stages: 50 pM, DE onward stages: 30 pM; probes are synthesized by Arbor Biosciences) in hybridization buffer (50% formamide, 10% dextran sulfate, and 5× Denhardt’s medium) was applied. Probes were denatured at 70°C for 10 min prior to mounting. Cellular DNA was co-denatured with the probes by placing the mounted coverslips directly on a heat source at 80°C for 10 min, followed by overnight hybridization at 37°C in a sealed, dark, humidified chamber.

Post-hybridization, coverslips were subjected to a stringency wash in pre-warmed 0.5× SSC containing 0.1% Tween-20 at 37°C for 30 min, followed by three 5-min washes in the same buffer at 42°C with shaking. Consecutive 10-min ambient washes were performed in 2× SSC with 0.1% Tween-20 and 1× SSC with 0.1% Tween-20. Following a final 1× PBS rinse, nuclei were counterstained with 3 μM DAPI in DPBS for 10 min at room temperature. Coverslips were mounted using VECTASHIELD Antifade Mounting Medium (Vector Laboratories, H-1000-10), sealed with nail polish, and stored at 4°C in the dark.

#### Confocal Imaging

For marker genes staining, images (1024×1024 pixels) were acquired in Galvano mode using a Nikon A1 confocal microscope (NIS-Elements v4.4) equipped with 405/488/561 nm lasers and an Apo TIRF 60× oil immersion objective (N.A. 1.49). Fluorescence was captured via GaAsP detectors (488/561 nm) or a MultiAlkali PMT (405 nm). Chromosome painting, speckle staining and histone modification imaging was performed on a Leica Stellaris 8 confocal laser scanning microscope (Leica Microsystems) using a HC PL APO CS2 100x/1.40 OIL objective.

Chromosome painting imaging acquired parameters: Excitation was done with the 405 nm laser and a white light laser tuned to 499 nm, and 653nm. Fluorescence emission was collected using detector ranges paired across three sequential acquisition passes to minimize spectral overlap as follows. Setting 1: HyD S 1(415-476nm) and HyD X 4 (665-752nm); Setting 2: HyD X 2(512-553nm). The confocal pinhole was set to 1.0 Airy Unit (152.7 um) for each channel. Images were acquired at a resolution of 512 x 512 pixels with a pixel size of 45nm and 3-line averages applied to reduce noise. Optical sections varying in total depth were taken where single z-stacks.

Histone modification imaging acquired parameters: Excitation was performed using a 405 nm laser line (for DAPI) and a white light laser tuned to 499 nm (for the 488 channel) and 590 nm (for the 594 channel). Fluorescence emission was collected across three sequential acquisition passes to minimize spectral overlap, configured as follows: Setting 1 (DAPI): HyD S 1 detector (415–476 nm); Setting 2 (488 channel): HyD X 2 detector (512–553 nm); Setting 3 (594 channel): HyD X 4 detector (605–660 nm). The confocal pinhole was set to 1.0 Airy Unit for each channel. Images were acquired at a resolution of 1024 x 1024 pixels with a pixel size of 45nm and 2-line averages applied to reduce noise.

Nuclear speckle related Imaging was performed on a Leica Stellaris 8 confocal laser scanning microscope (Leica Microsystems) equipped with an HC PL APO CS2 100x/1.40 OIL objective. Excitation was performed using a 405 nm laser (2.30% power) and a white light laser tuned to 499 nm (6.31%), 579 nm (5.77%), and 653 nm (15.35%). Fluorescence emission was collected using sequential acquisition passes to minimize spectral overlap as follows. Pass 1: HyD S Channel 1 (425–430 nm) and HyD X Channel 4 (601–829 nm); Pass 2: HyD X Channel 2 (504–595 nm) and HyD S Channel 3 (430–492 nm). The confocal pinhole was set to 1.0 Airy Unit (130.9 µm) for each channel. Images were acquired at a frame size of 432 × 432 pixels with a pixel dwell time of 1.567 µs and 2-line averaging applied to reduce noise. Optical z-sections were captured at a z-step size of 0.3 µm. Real-time computational deconvolution was performed using the Leica LIGHTNING module within Leica Application Suite X (LAS X) to enhance spatial resolution and signal-to-noise ratio.

All acquisition parameters were controlled using Leica Application Suite X (LAS X 4.6.1.27508).

### Image analysis and quantification

#### Chromosome painting image analysis

Exported DNA FISH confocal images formatted as “.tiff” were analyzed using a custom Python pipeline implemented in a Jupyter Notebook (https://github.com/dekkerlab/4DN-Hardening-paper) to quantify chromosome territory morphology and spatial organization. Nuclear regions were segmented from the DAPI channel using Gaussian smoothing, Otsu thresholding, and morphological filtering to generate binary nuclear masks. Images without a successfully segmented nucleus were excluded from subsequent analyses.

Chromosome territories were identified from the chromosome painting channels corresponding to chromosome 15 (488 nm) and chromosome 11 (550 nm). Within each nucleus, chromosome territory masks were generated using Gaussian smoothing followed by stage-specific adaptive intensity thresholding to account for differences in fluorescence intensity across differentiation stages. Binary masks were refined by morphological opening and closing, removal of small objects, and hole filling to eliminate non-specific signals. When multiple connected components were detected within a chromosome mask, an adaptive connected-component algorithm was used to distinguish separated homologous chromosome territories from fragmented signals based on the relative size and spatial separation of connected objects.

The boundaries of the nucleus and chromosome territories were extracted from the binary masks using OpenCV contour detection, and centroid coordinates were recorded for each segmented object. Convex hulls were computed from the extracted contours to define the outer boundary of each chromosome territory. Chromosome territory morphology was quantified by measuring territory area, perimeter, perimeter-to-area ratio, and normalized radius of gyration. Spatial overlap between chromosome 11 and chromosome 15 territories was quantified using both the overlap area and the Jaccard index. Measurements from all successfully segmented nuclei were combined for downstream statistical analyses and visualization.

#### Immunofluorescence

Exported immunofluorescence confocal images formatted as “.tiff” were analyzed using a custom Python pipeline implemented in a Jupyter Notebook (https://github.com/dekkerlab/4DN-Hardening-paper). Nuclear masks were generated from the DAPI channel using Gaussian smoothing, Otsu thresholding, and morphological opening and closing operations. Small objects and holes were removed, and the largest connected component was retained as the nuclear mask. Images without a successfully segmented nucleus were excluded from further analysis.

Histone modification signals (H3K9me2, H3K9me3, or H3K27ac) were extracted from the corresponding fluorescence channel and smoothed using a Gaussian filter before quantification. Mean nuclear fluorescence intensity was calculated as the average pixel intensity within the nuclear mask, and the integrated nuclear intensity was obtained by summing all pixel intensities within the nucleus.

High-intensity histone domains were identified within each nucleus by applying additional Gaussian smoothing followed by thresholding at the 80th percentile of nuclear fluorescence intensity. The resulting binary masks were refined by morphological opening and closing, removal of small objects, and hole filling to eliminate isolated noise. For each nucleus, the total area, mean fluorescence intensity, and integrated fluorescence intensity of high-intensity domains were quantified. The radial localization of high-intensity domains was calculated as the intensity-weighted mean normalized distance of domain pixels from the nuclear centroid, where distances were normalized by the average nuclear radius derived from the nuclear boundary. Measurements from all successfully segmented nuclei were pooled for downstream statistical analyses and visualization.

### 3D volume analysis

Representative images were generated by Fiji. As SON intensity varied across images of the same treatment, image stacks were first normalized for intensity using a custom analysis (https://github.com/dekkerlab/4DN-Hardening-paper/Fig4_EDFig5_SpeckleVolume). Briefly, a single global normalization factor for the SRRM2 channel was computed for each stack. We then divide every voxel by this factor to preserve the relative intensity differences between z-slices. These normalized images were then saved as tif files and imported into Leica-compatible Aivia software (https://www.aivia-software.com) for 3D segmentation and volume analysis using the “3D object analysis” tool. Speckle volume and speckle count per cell data are then extracted from the Aivia output and plotted.

### RNA-seq analysis

Reads from both experiments (H1-derived five-stage series with 2 biological replicates per stage, and H9-derived DE-to-HB eight-day series) were adapter-trimmed with fastp and quantified with Salmon (v1.10.3) against a decoy-aware GENCODE v50 index. Transcript-level estimates were imported and summarized to genes with tximport (v1.34.0) using countsFromAbundance=“no” to pass average transcript lengths to DESeq2 so that changes in isoform composition are accounted for within the model. Mitochondrial genes were removed before normalization, and nuclear genes with at least 10 counts in at least 2 samples were retained. We derived expression tables from the DESeq2 (v1.46.0), including TPM, variance-stabilized transformed (VST) values, as well as GeTMM (edgeR v4.4.0), which is normalized for both gene length and library composition for comparisons spanning genes and samples at once. Differential expression was run separately for the two experiments using DESeq2. For the five-stage series, the four sequential transitions were tested independently (DE vs ESC, HB vs DE, iHLC vs HB, HLC vs iHLC) with Wald tests and apeglm log2-fold-change shrinkage. A likelihood-ratio test (LRT) was also run across the five stages. For the eight-day series, a continuous design of expression as a function of the timepoint (day) was tested against an intercept-only model by LRT.

### Gene set enrichment analysis

Preranked gene-set enrichment analysis was performed with gseapy (v1.1.13), ranking all genes in a transition by their DESeq2 Wald statistic for the four sequential transitions with 1,000 permutations, gene-set size 5-1,000 and a fixed seed. Two gene-set resources were used: the 50 MSigDB Hallmark gene sets (release 2024.1.Hs) and the Enrichr KEGG pathway library (KEGG_2026). Gene sets were called enriched at FDR q < 0.05 in at least one transition. For display, Hallmark sets were grouped by their MSigDB process category, and KEGG pathways were annotated with the KEGG BRITE hierarchy (br08901) with only the Environmental Information Processing and Cellular Processes categories shown.

### ATAC-seq analysis

ATAC-seq libraries were processed with the ENCODE-DCC ATAC-seq pipeline (v2.2.3). For downstream analysis, Tn5 insertion positions were extracted from the filtered fragments with a 9-bp offset correction. Rather than defining a consensus region set from peak calls, insertion positions were quantified over the ENCODE registry of candidate cis-regulatory elements (cCREs) version 4 for GRCh38. Tn5 insertion counts for cCREs were normalized to log2 counts-per-million, and differentially accessible cCREs were identified for each of the four stage transitions using a limma-trend moderated-variance model ^68^. Among elements with detectable accessibility (CPM > 1) in at least one of two compared stages, elements were called differentially accessible at greater than 2-fold change and a Benjamini-Hochberg FDR < 0.05.

The union of differentially accessible cCREs across all transitions was partitioned into accessibility archetypes by fuzzy c-means clustering of the five-stage accessibility trajectory (per-stage mean normalized log2-CPM, z-scored by stage, fuzzifier m=2). The number of clusters (C=8) was selected as a local minimum in the Xie-Beni index in a sweep of C=4-10. For display, cCREs within each of the archetypes were sub-clustered using EVoC (0.3.1) on the four marks H3K4me3, H3K27ac, H3K27me3, H3K9me3 across the 5 stages (20 features, mean fold-change over input over each cCRE z-scored), with sub-clusters smaller than 400 cCREs folded into a single “diffuse” group. Dendrograms between the primary sub-clusters were calculated by Ward linkage clustering on their centroids.

### TF motif enrichment at differentially accessible cCREs

TF motif enrichment was computed for the differentially accessible cCREs at each stage transition with AME (MEME Suite v5.5.9), using the up- and down-accessibility cCREs of each of the four sequential transitions (eight sets in total) as foreground sets. For each foreground set, a GC-content-matched background was constructed by sampling five times as many regions from expression-filtered cCREs excluding the foreground, matched to the foreground GC distribution in 2% bins. Motifs were tested with AME using average odds scoring and Fisher’s exact test against the matched background, and a log2 odds-ratio was computed for each motif from the true/false-positive counts, using pseudocounts of 0.5. The motif panel was a curated set of 91 position-weight matrices from distinct TFs drawn from HOCOMOCO (H14CORE), plus the POU5F1::SOX2 composite motif from JASPAR. These were grouped into thirteen TF families spanning the trajectory’s regulators. In each contrast, motif enrichment was integrated with driver TF expression. Taking the “active” stage to be that in which the foreground cCREs are accessible, family members were required to be expressed in the active stage (TPM ≥ 1) and to have a significant, non-trivial motif enrichment (AME adjusted p < 0.05 and log2 odds-ratio ≥ 0.5). The top three surviving factors were displayed, ranked by expression.

### CUT&RUN and ChIP-seq analysis

Chromatin-profiling libraries (CUT&RUN for H3K4me3, H3K4me1, H3K27me3, H3K36me3, H3K9me2, H3K9me3 H4K20me3 and ChIP-seq for H3K27ac) were processed with the ENCODE-DCC ChIP-seq pipeline (chip-seq-pipeline2, v2.2.2). Histone modifications were processed in “histone” mode, each against its matched input (ChIP-seq) or IgG (CUT&RUN) control. Genome-wide signal tracks were generated as MACS2 fold-enrichment-over-control coverage and peaks were called under the pipeline defaults for the respective mode.

### Hi-C analysis

Hi-C data were processed with distiller-nf (v0.3.4). Reads were mapped with *bwa mem* to the GRCh38 no-alt analysis set including the hs38d1 decoy, and ligation junctions were classified with *pairtools* and duplicates were removed with *pairtools*. Valid pairs were binned into multi-resolution cooler (.mcool) matrices after filtering (MAPQ ≥ 30). Matrices were balanced by iterative correction (*cooler balance*) genome wide. Contact matrix features were computed with cooltools (v0.5.1 or later), bioframe (v0.8.0) and jointly-hic (v1.0.0 or later). To generate genome-wide P(s) curves, *cis* contact frequencies were computed from balanced contact maps separately for each chromosome arm at 1-kb resolution, followed by aggregation across chromosome arms and smoothing of the expected contact frequencies (Gaussian smoothing parameter, σ = 0.01).

For the five-stage experiments, compartmentalization was quantified at 50-kb resolution. Rather than each stage’s own eigenvector, bins were ranked by the scores of the leading joint interaction-profile principal component PC1 (see below), which provides a common compartmentalization axis across all five stages. Observed contact frequencies were divided by the distance-decay expected, averaged within a grid of 38 quantile bins of PC1 score, after excluding the upper and lower 2% tails, and average observed/expected contact frequencies were calculated between all bin pairs to generate compartment saddle matrices. For the eight-day experiments, compartmentalization was quantified by *cis* eigenvector decomposition of balanced Hi-C contact matrices at 100-kb resolution using *cooltools eigs-cis*, using a hg38 chromosome-arm view and GC-content as the covariate for eigenvector phasing.

Chromosome arm-specific expected cis contact frequencies were calculated using cooltools *expected-cis* and saddle plots were then generated using *cooltools saddle*, in which genomic bins were grouped by E1 rank into 38 equally populated bins, after excluding the upper and lower 2.5% tails, and average observed/expected contact frequencies were calculated between all bin pairs to generate compartment saddle matrices. For both experiments, compartment strength was quantified using the cooltools saddle-strength metric, defined as log2[(AA + BB) / (AB + BA)] from the four corners of the saddle matrix as a function of the quantile threshold defining those corners.

Insulation scores for the eight-day experiment were calculated from balanced Hi-C contact matrices using the *cooltools.insulation* API. Diamond insulation scores were computed at 10-kb resolution using a 100-kb sliding diamond window, and boundary strengths were identified using the Li thresholding method implemented in cooltools. Aggregate insulation profiles were generated by centering all called insulation boundaries on their midpoint and averaging insulation scores across a ±100-kb genomic window surrounding each boundary. Aggregate profiles were used to compare insulation patterns across different days in DE to HB transition. Insulation strength was quantified as the difference between the maximum and minimum values of the aggregate insulation profile for each sample, with larger values indicating stronger insulation at domain boundaries.

Chromatin loop analysis across the eight-day experiment was performed using cooltools. A common set of chromatin loops identified from the HB Hi-C dataset was used for all analyses. Balanced 5-kb Hi-C contact matrices were first normalized by chromosome arm-specific expected cis contact frequencies. Aggregate loop pileups were then generated using *cooltools.pileup*, centered on the intersection of loop anchors with a ±100-kb flanking region around each loop. Pileup matrices from all loops were averaged to generate a single observed/expected aggregate loop interaction matrix for each sample. Aggregate loop strength was quantified from the averaged pileup matrix as the enrichment of the central loop pixel relative to the local background. The background interaction frequency was calculated as the mean signal from the four corner regions and four adjacent stripe regions surrounding the central pixel, and loop strength was defined as the ratio of the central pixel intensity to the average background signal. Aggregate pileup matrices were visualized as log2(observed/expected) interaction maps, and loop strength values were compared across differentiation stages.

### IPT annotation and characterization

To localize changes in long-range folding along the genome in a comparative manner, the autosomal contact maps of all five stages were jointly decomposed by their *trans* (inter-chromosomal) interaction profiles at 50-kb resolution using the *jointly-hic* framework ^42^. For each stage’s balanced whole-genome map, the trans interaction profile of every 50-kb bin was extracted following preprocessing (masking of low-coverage/outlier bins, imputation of *cis* blocks from sampled trans contacts, and iterative correction to a doubly-stochastic affinity matrix). A single incremental principal component (PC) model (32 components) was trained across all five stage maps so that they share one set of basis vectors, and each stage was then projected onto this shared basis, yielding, for every 50-kb bin, a profile of principal-component scores across the five stages.

To resolve the temporal behavior of each locus, bins were clustered on the full cross-stage score vectors using the leading six components across all five stages by k-means (k-means++ initialization, 100 restarts). A ten-cluster solution was retained and manually consolidated to form seven Interaction Profile Trajectories (IPTs) by merging pairs of clusters that differed only in centromeric-versus-telomeric proximity of otherwise identical profiles. The seven-trajectory annotation was used for all downstream epigenetic, saddle and gene-level analyses.

Contact enrichment between the seven interaction-profile trajectories was summarized as a “discrete saddle”: for every pair of IPT categories, observed contacts were divided by the distance-decay expected and averaged over all bin pairs in each category combination, excluding the first two diagonals, at 50-kb resolution over the autosomes. Enrichment was computed for *cis* and *trans* contacts and displayed as log2(mean observed/expected).

To relate the interaction-profile trajectories to independent maps of nuclear landmark association, published domain annotations were overlaid onto the 50-kb IPT bins. LAD calls were taken from public 4DN LMNB1 DamID/pA-DamID accessions for seven cell lines, including an H1-hESC map and six other cell lines (4DNFIJXADI29, HFFc6: 4DNFIT9W77EE, K562: 4DNFIJHD22QE, HCT 116: 4DNFIA2LBQCD, HAP1: 4DNFIDFCY3JN, RPE-hTERT: 4DNFIUVTO2H3, U2OS: 4DNFIZ3JHKWC) and total coverage of LAD annotations was calculated per IPT. SPIN states were overlaid from K562 (Wang et al. 2021) and from H1 cells from 4DN (4DN WA01 p30 WB35186 (human H1ESC)). Enrichment was calculated as the fraction of IPT bins per SPIN state.

### IPT expression trajectories and cell-type lineage enrichment

Genes were assigned to 50-kb IPT bins based on the transcription start site. Within each IPT, genes were split into a down-regulated and an up-regulated set at the DE-to-HB step based on apeglm-shrunken HB-vs-DE absolute fold-change > 2 and cross-stage LRT padj < 0.05. For each set, the five-stage VST trajectory was averaged across genes and plotted as mean ± SEM at each stage. Within each IPT, genes were split into a down-regulated and an up-regulated set at the DE-to-HB step. For each set, the five-stage variance-stabilized expression trajectory (per-stage mean of replicates) was averaged across genes and plotted as mean ± SEM at each stage. For each IPT, preranked gene-set enrichment (gseapy prerank) was run over that trajectory’s genes ranked by their HB-vs-DE DESeq2 Wald statistic, tested against PanglaoDB cell-type marker sets obtained from Enrichr (PanglaoDB_Augmented_2021).

### Promoter and cCRE clustering

H3K27me3 and H3K27ac fold change signal was quantified in ±5 kb windows centred on T.pcg gene promoters or on T.pcg cCRE midpoints. Each element was represented by the mean signal over the central bins of both marks across the five stages, which were log-transformed, standardized, and partitioned by k-means. All T.pcg promoters were clustered, while for cCREs, 5,000 T.pcg cCREs were sampled at random. Stackup heatmaps of bigWig signals were generated using pybigtools and matplotlib.

### Average inter-arm interactions

Average inter-arm interactions were quantified from MAPQ 30-filtered Hi-C matrices with merged replicates at 100-kb resolution for five differentiation stages (ESC, DE, HB, iHLC and HLC). Analyses were restricted to the 22 autosomes, giving 231 unique interchromosomal chromosome pairs. Bins overlapping the predefined low-quality-bin mask were excluded. In addition, the terminal 1 Mb of each chromosome arm and the 1-Mb regions adjacent to centromeres were masked to reduce mapping artefacts.

For each chromosome pair, the balanced contact matrix was divided into the four possible chromosome-arm combinations: p–p, p–q, q–p and q–q. Expected contact frequency was defined as the mean balanced interchromosomal contact frequency across the corresponding whole-chromosome pair. To retain zero-valued pixels, a chromosome-pair-specific pseudocount, p, was defined as the smallest finite positive contact value among the four arm-pair matrices. The same value was added to both observed and expected contacts and observed/expected enrichment was calculated as (O+p)/(E+p).

For heatmap visualization, each arm-pair O/E matrix was linearly rescaled to 100 × 100 bins so that chromosome arms contributed equally irrespective of their genomic length. The four rescaled matrices were assembled into a 200 × 200 map, with the centromere-centromere position at the center and telomere-telomere positions at the four outer corners. Chromosome-pair matrices were averaged with equal weight within each differentiation stage and symmetrized by averaging each matrix together with its transpose. Iterative correction was applied after the chromosome-pair maps had been averaged. Correction was performed on the finite, non-empty portion of the averaged matrix, using a maximum of 100 iterations and a convergence tolerance of 10^{-5}. The corrected matrices were log2-transformed for plotting. Heatmap axes represent distance from the centromere as chromosome-arm percentiles, ranging from the centromere at 0% to the telomeres at 100%. Note that iteratively corrected matrices were not used for score or decay calculations.

### Centromere-centromere and telomere-telomere interaction scores

Interaction scores were calculated separately for every chromosome pair from the uncorrected O/E matrices. O/E values were first log2-transformed and then averaged across the relevant pixels. Thus, each score represents the mean log2 O/E, rather than the log2 transformation of an arithmetic mean. Scores were calculated within 4 Mb of the corresponding telomere/centromere along each chromosome axis: the central centromeric window or the four terminal telomeric windows. All centromeric pixels were averaged per chromosome to produce a single score for each chromosome. The same procedure was repeated for telomeric pixels. This produced one centromeric and one telomeric score per chromosome pair and stage.

For the five-stage differentiation trajectory from H1 hESCs to HLCs, chromosome-pair scores were visualized as violin plots with an internal boxplot. The zero line indicates equal observed and expected contact frequency. Chromosomes were divided using the operational classification in the analysis notebook: chromosomes 1–10 were designated “large” and chromosomes 11–22 “small.” Adjacent differentiation stages were compared using Wilcoxon signed-rank tests, with scores matched by chromosome pair across stages. The implemented test selected the one-sided alternative from the sign of the median paired difference. P-values were adjusted within each family of comparisons using the Benjamini–Hochberg false-discovery-rate procedure.

For 8-day DE-to-HB transition, chromosome-pair scores were visualized as line plots. Thin lines show individual chromosome trajectories, while the prominent line shows the mean across chromosomes at each stage. Shading indicates the interquartile range. Ordered trends were assessed with chromosome-blocked Page tests, specifying a decreasing trend for centromeric scores and an increasing trend for telomeric scores. Plots report the Page statistic, the unadjusted P value, and the change in mean score between DE and HB.

Centromeric and telomeric trans-contact decay profiles were derived from the unrescaled, uncorrected arm-pair O/E matrices (100 Kb for telomeres, 250 Kb for centromeres). For each chromosome-arm pair, pixels were grouped according to their summed two-locus distance from the centromere or telomere. For example, a summed distance of 10 Mb represents approximately 5 Mb from the landmark at each locus when the two offsets are equal. Distances up to 15 Mb were retained. At each distance, contact values were obtained along the corresponding diagonal extending from the centromeric or telomeric corner of each arm-pair matrix. Diagonals were center-cropped to a maximum of 16 pixels, corresponding to the configured 4-Mb diagonal width at 250-kb resolution. Positive finite O/E values were log2-transformed and averaged to obtain one value for each chromosome-arm pair and distance. Curves show the arithmetic mean of these arm-pair log2 O/E values, with shaded two-sided 95% bootstrap confidence intervals calculated using 1,000 resamples and a fixed random seed of 0. A distance was displayed only when at least five chromosome-arm pairs contributed.

### DpnII-seq analysis

DpnII-seq libraries were processed using https://github.com/dlafont15/DpnII-seq. Briefly, sequenced reads were mapped to hg38 using the bowtie read aligner ^69^ and reads mapping to multiple regions of the genome were removed. To remove artificial biases, we filtered out paired-end reads from fragments where cut-sites were more than three nucleotides away from the start or end of the paired-end reads. Filtered reads were then binned at a resolution of choice. To account for copy number variation due to karyotype and rearrangements in given cell lines, we leveraged publicly available copy number data from the CatalOgue of Somatic Mutations In Cancer (COSMIC) database to assign a copy number state to each genomic bin. Read coverage files were corrected to a genome-wide diploid state using the copy number assignments and dividing coverage by an appropriate correction value (i.e. diploid = 1, triploid = 1.5, tetraploid =2, etc). Since odd fluctuations were not observed after plotting DpnII-seq along chromosomes and that ESC cell lines are largely assumed to be diploid, we set a genome-wide scalar of 1. Final copy number corrected coverage files were used for all downstream analyses.

### Liquid Chromatin Hi-C analysis

The analysis of LC-Hi-C in the is paper largely builds from the work done in ^48^ and was written in R and python as both executables and scripts to be run R Studio and bash, respectively. All scripts can be found at “https://github.com/dekkerlab/4DN-Hardening-paper/LCHiC/” which some functions were adapted from https://github.com/tborrman/liquid-chromatin-Hi-C/tree/master/src/scripts. The Loss of structure (LOS) metric quantifies the loss of interactions within a 2Mb window along the diagonal after DpnII pre-digestion at every bin ^48^. This is accomplished by first calculating the percent cis interactions within a 2Mb window and dividing this by the total interactions (cis and trans) within that bin for both the mock- and DpnII-digested Hi-C libraries. LOS is then calculated by obtaining the difference between the percent cis values and normalizing by the mock-digested percent cis values:

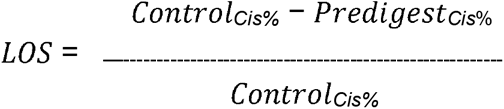

Note that interactions with a span less than 1kb were discarded in the cooler files used to generate cispercent.bedgraph files.

The DpnII-seq sample is used to normalize our LOS metric. This is done by first plotting LOS against DpnII-seq signal and calculating a moving average across the data. LOS residuals are then determined by calculating the difference between each data point and the moving average.

### Fragment size analysis

Since functionality for scaling and plotting fragment size distribution curves in Agilent PROsize software is limited, we employ a custom R script to calibrate and normalize the migration time over fluorescence output from Agilent Fragment Analyzer for plotting. Briefly, for each sample of a Fragment analyzer run, migration time over relative fluorescence data is extracted from ‘.raw’ files manually using PROsize software. For each Fragment Analyzer run, the migration time of each peak in the standard ladder is also obtained manually from the .raw files using the PROsize GUI. Piecewise linear functions are then calculated between each peak in the ladder to predict size (in bp) from migration time for specific migration time intervals. The linear models generated from the ladders are then used to convert migration times into fragment sizes for each sample of that run. To compare samples between runs, ladder migration times for each peak across the ladders (of each run to be compared) are averaged to generate an averaged calibration curve. Piecewise functions are again generated between each peak of the averaged calibration ladder, but this time to convert predicted size to a common normalized time axis so we can directly plot and compare samples from different runs. Relative fluorescence is then scaled to the highest non-marker peak which is most often the maxima generated by DpnII digestion. Upper and lower bounds are then scaled to the upper and lower markers, respectively, of the Agilent sample loading buffer for HS Large Fragment kits. Scaled time curves are then plotted and the average migration time for each peak in the averaged calibration ladder are used to label and tick the x-axis with fragment size values.

## Data availability

Any requests regarding cell lines should be directed to R.M. and J.D. The datasets generated in this publication have been deposited in the NCBI GEO as a SuperSeries accessible through the accession numbers, consisting of GSE343495 (bulk Hi-C & Liquid Hi-C), GSE343496 (RNA-seq), GSE343493 (ChIP-seq), GSE343494 (CUT&Run) and GSE343497 (ATAC-seq). All other data supporting the findings of this study are available from the corresponding author on reasonable request.

## Code availability

Custom scripts and notebooks used in this study are publicly available on GitHub: https://github.com/dekkerlab/4DN-Hardening-paper/, and https://github.com/abdenlab/paper-4dn-stepwise.

## Acknowledgements

We thank all the members of the Dekker lab, Abdennur lab, Maehr lab and Mirny lab and for helpful discussions. We thank all members of 4DN Center for 3D Structure and Physics of the Genome. We thank Core Facilities at the University of Massachusetts Chan Medical School for their assistance: the Deep Sequencing Core, the Flow Cytometry Core, and Sanderson Center for Optical Experimentation (SCOPE) Core. Imaging data was acquired with support from C. Baer and J. McConnell on the Leica STELLARIS 8 STED confocal microscope. Diagrams included in figures were created with BioRender.com.

## Funding

Imaging analysis using the Aivia software on the Aivia workstation was funded by a Massachusetts Life Science Center Research Infrastructure grant award to Drs. K. Fitzgerald and C. Baer. The SCOPE RRID is SCR_022721. This work is supported by the 4DN Center for 3D Structure and Physics of the Genome, funded by the National Institutes of Health Common Fund: grant HG011536 to J.D. and L.M., and the National Human Genome Research Institute: grant HG011583 to D.L.L., and the NIH National Center for Advancing Translational Sciences: grant T32TR005445. J.D. is an investigator of the Howard Hughes Medical Institute.

## Author Contributions

Conceptualization: J.H.G., N.A., L.M., J.D., R.M., X.H., D.L.L., T.M.R.; Differentiation: R.S., J.H., J.M., K.M.P., X.H., L.Y.; Hi-C experiments: L.Y.; ChIP-seq experiments: L.Y.; CUT&RUN-seq experiments: J.H., J.M., K.M.P.; LC-Hi-C experiments and analysis: D.L.L., L.Y.; Imaging and Analysis: X.H., D.L.L.; 5-stage RNA-seq, ATAC-seq, ChIP-seq, CUT&RUN, and Hi-C processing and analysis: N.A., T.M.R., V.Y., H.T.; IPT analysis: T.M.R., N.A.; DE-to-HB 8-day transition experiments: L.Y., and J.H.G.; DE-to-HB 8-day transition analysis: J.H.G., H.T., N.A., T.M.R., X.H.; Telomeric and centromeric analysis: A.A.G.; T.pcg analysis: N.A., E.N.; Writing – original draft: N.A., T.M.R., J.H.G., X.H., D.L.L., J.D.; Writing – review and editing: N.A., E.N. X.H., D.L.L.,J.H.G., L.M., R.M., J.D.; Visualization: N.A., X.H., D.L.L.; Supervision: J.H.G., N.A., L.M., J.D.; Funding acquisition: J.D., L.M., J.H.G., N.A., R.M., D.L.L.

## Conflicts of interest

Job Dekker is a member of the scientific advisory board of Arima Genomics, San Diego, CA, USA.

## Extended Data Figure

**ED Fig 1:**
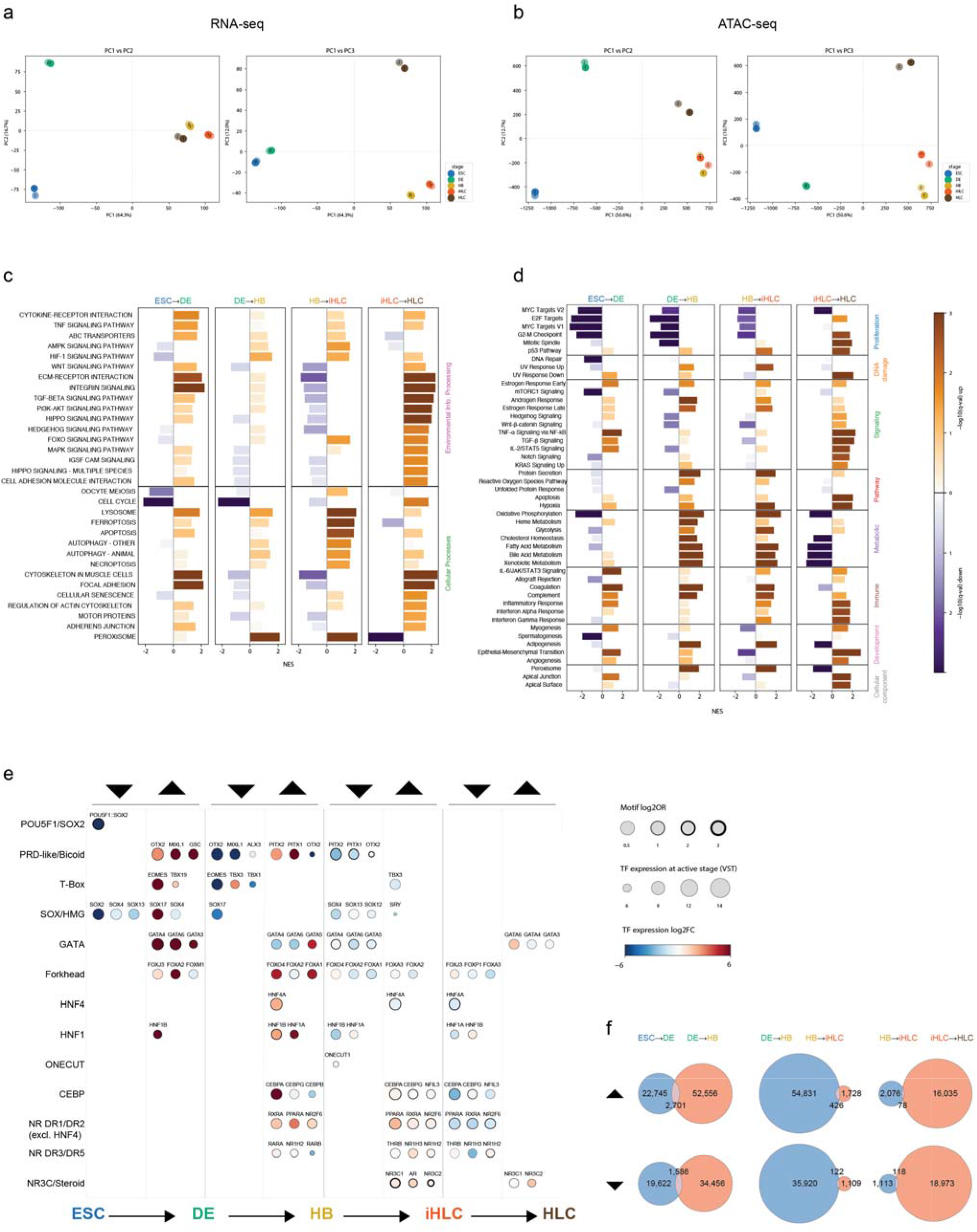
Replicate, gene set enrichment and TF-motif enrichment analyses. Principal component analysis of bulk RNA-seq (**a**) and ATAC-seq replicates (**b**). Gene set enrichment analysis was performed using either KEGG (**c**; KEGG database) or Hallmark (**d**; MSigDB) gene set collections. Normalized Enrichment Scores (NES) are presented for each grouping. **e**, Transcription factor (TF) motif-enrichment analysis integrating chromatin accessibility and TF expression data at each transition. Each transition is subdivided into significantly increased and reduced accessibility contrasts. Bubbles depict the top 3 or fewer candidate driver TFs for the corresponding contrast of differentially accessible elements. Bubble size reflects expression of the TF in the “active” stage of the contrast where the elements have greater accessibility. Bubble color reflects the fold change in expression of the TF across the transition. Bubble border size reflects the TF motif log odds ratio for the given contrast. **f**, Venn diagrams showing overlap of upregulated (top row) and downregulated (bottom row) regulatory elements between successive transition steps.

**ED Fig. 2:**
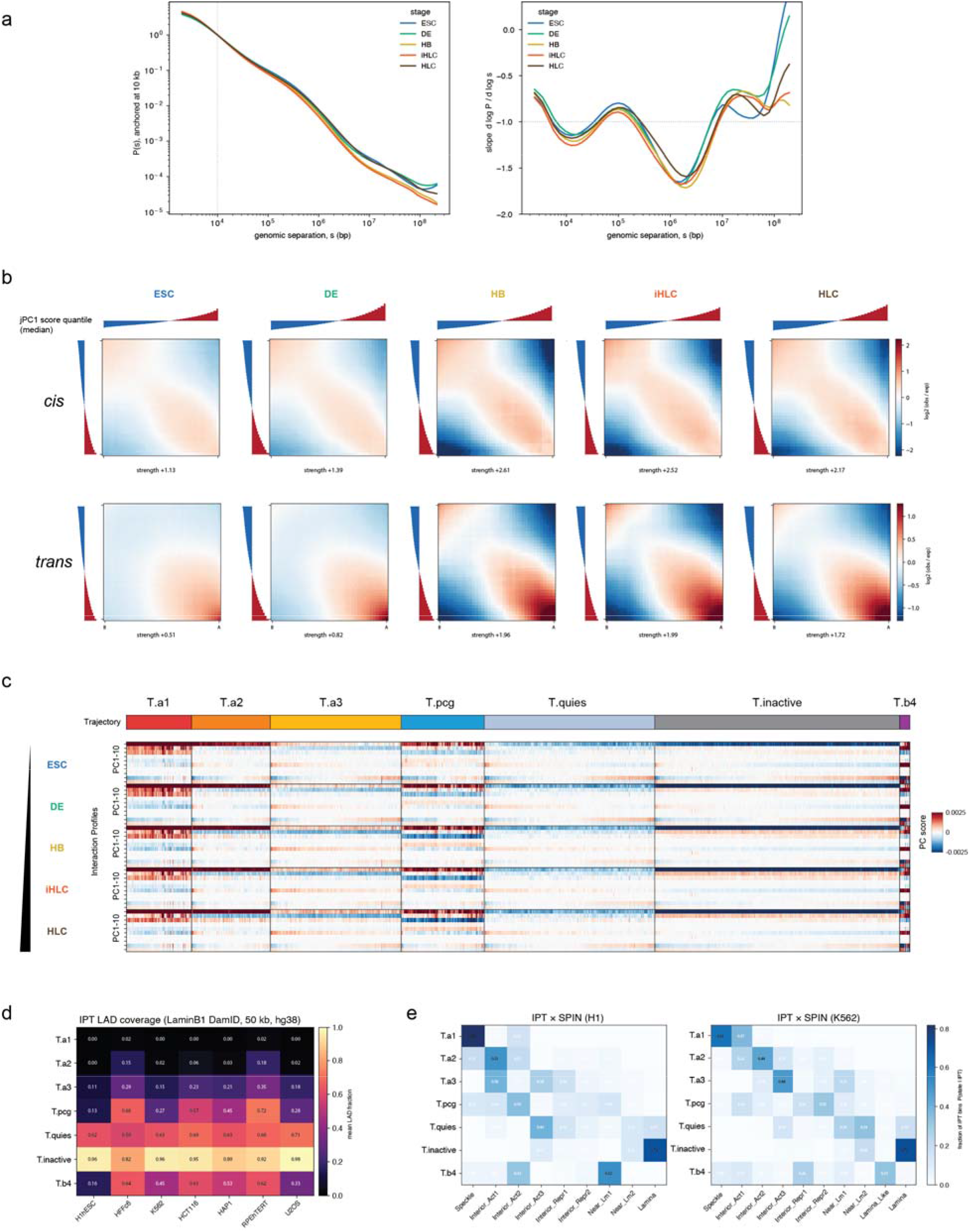
Hi-C analysis, joint-PCA scores, and LAD and SPIN state characterization of IPTs. **a**, P(s) plots for Hi-C data from each stage during in vitro hepatic differentiation and associated derivative plots. **b**,Saddle plots of *cis* (top) and *trans* (bottom) contact enrichment, ranking bins by their jointly derived *trans* PC1 score vectors via joint PCA. Saddle strength scores are calculated using 20% corner extents of the saddle matrix. **c**, Heatmap of joint PC1–PC10 score vectors for each stage. Columns represent 50-kb bins, grouped by IPT cluster (top row), and ordered as in Fig. 2b. **d**, Fraction of each IPT covered by LAD annotations generated from Lamin DamID data for 7 different cell lines. **e**, Correspondence between SPIN and IPT annotations for H1 hESC (left) and K562 (right) cell lines. Color scale indicates the fraction of IPT bins assigned to each SPIN state, with rows summing to 1.

**ED Fig. 3:**
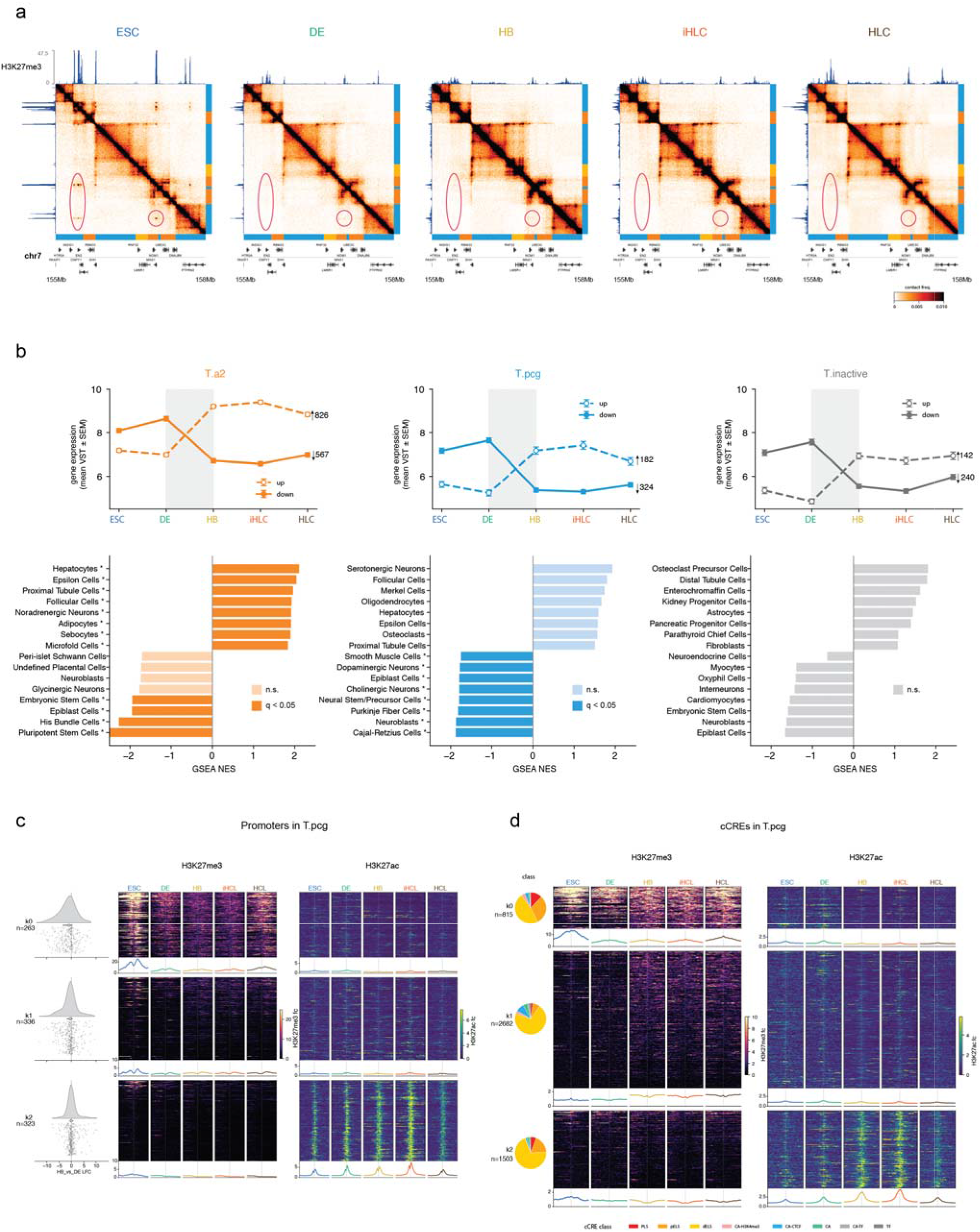
The T.pcg trajectory captures H3K27me3 spreading and lineage-specific gene restriction at the DE-to-HB transition. Hi-C contact frequency maps of a region on chromosome 7 highlighting changes in interaction frequency and H3K27me3 distribution. Regions containing sharp, punctate polycomb interactions in hESC that vanish in DE onward are circled. This coincides with reduction in H3K27me3 peaks in DE, followed by spreading across T.pcg domains in HB onward. **b**, Top: Mean expression trajectories for genes significantly up- and down-regulated at the DE-HB transition, restricted to T.a1 (left), T.pcg (middle), and T.inactive (right) regions of the genome. Error bars depict standard error. Bottom: Gene Set Enrichment Analysis of all differentially expressed genes at the DE-HB transition against the PanglaoDB cell-type gene marker sets, restricted to T.a1 (left), T.pcg (middle), and T.inactive (right) regions of the genome. The top and bottom 8 genes by Normalized Enrichment Scores (NES) are shown. Dark-colored bars depict hits at 0.05 FDR; light-colored bars are below significance. **c,** Stackup heatmaps of H3K27me3 and H3K27ac histone modification signal centered on TSSs in T.pcg, grouped by jointly clustering by signals across the five stages. Kernel density plots and point scatters on the left depict the distributions of log fold change in expression of the corresponding genes in each cluster. Cluster blocks are sorted by descending mean H3K27me3 at HB over a central window of 600 bp, while rows within each cluster block are sorted by descending H3K27me3 averaged across all stages over the same window. **d,** Stackup heatmaps of H3K27me3 and H3K27ac histone modification signal, as in **c**, but centered on 5,000 randomly sampled cCREs in T.pcg. Pie charts on the left depict the cCRE class distribution corresponding to each of the three clusters.

**ED Fig. 4:**
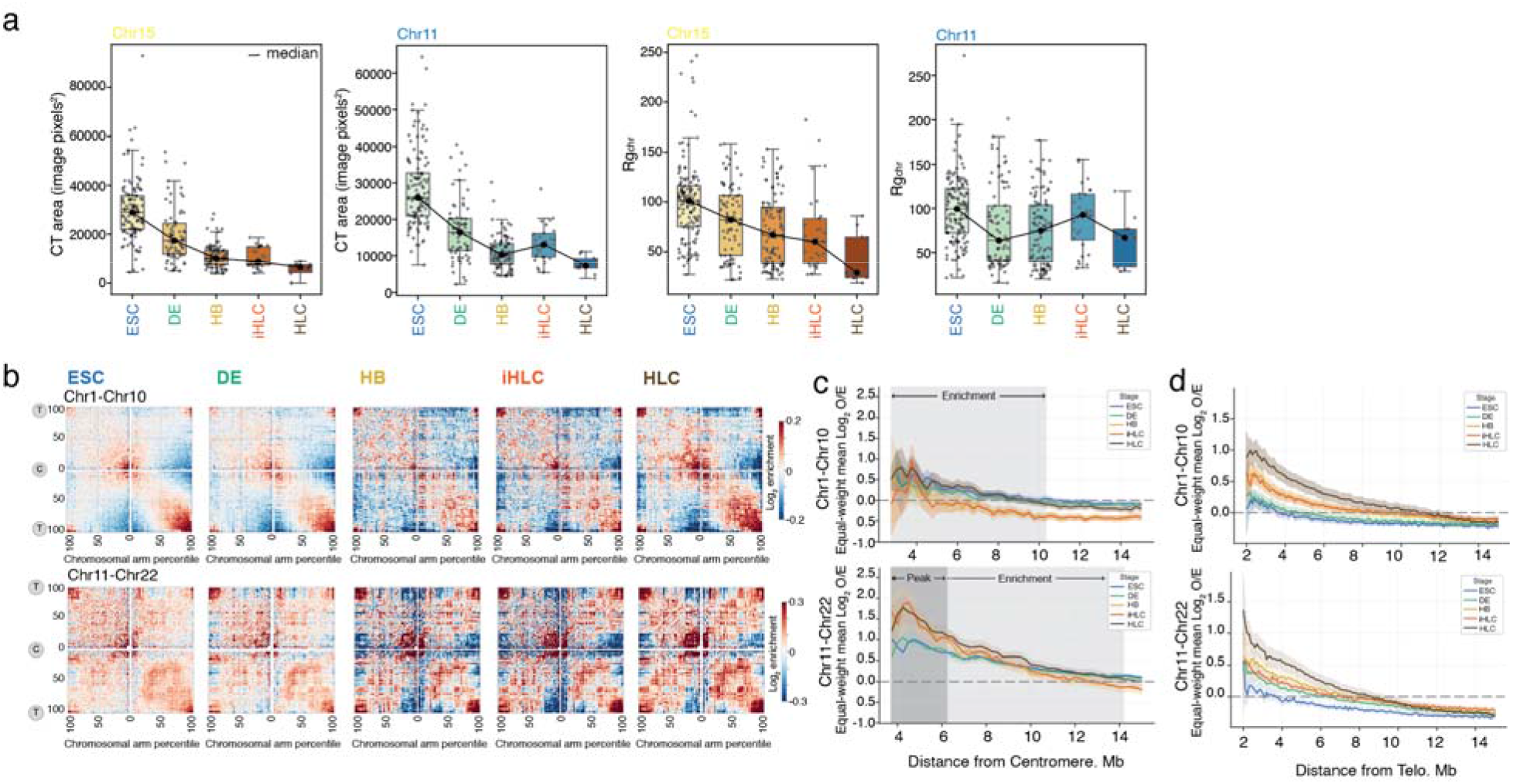
Centromeric and telomeric interaction dynamics across regions and different size of chromosomes. **a**, Quantification of territorial properties of chromosomes 11 and 15 measured from chromosome painting across five stages. Left: pixel images of chromosome territory (CT) area; right: radius of gyration of each CT. **b**, Scaled-chromosome saddle plots showing piled-up interchromosomal interactions of large (chr1–chr10) or small (chr11–chr22) chromosomes across five stages. “T” represents telomeric regions; “C” represents centromeric regions. **c-d**, Quantifying decay of average interaction strength at pericentromeric (**c**) and telomeric (**d**) regions between pairs of large chromosomes (top) and pairs of short/small chromosomes (bottom). Shaded areas show visible enrichment of pericentromeric interactions above expected for large and small chromosomes (c) and a relative peak at shorter distances for small chromosomes.

**ED Fig. 5:**
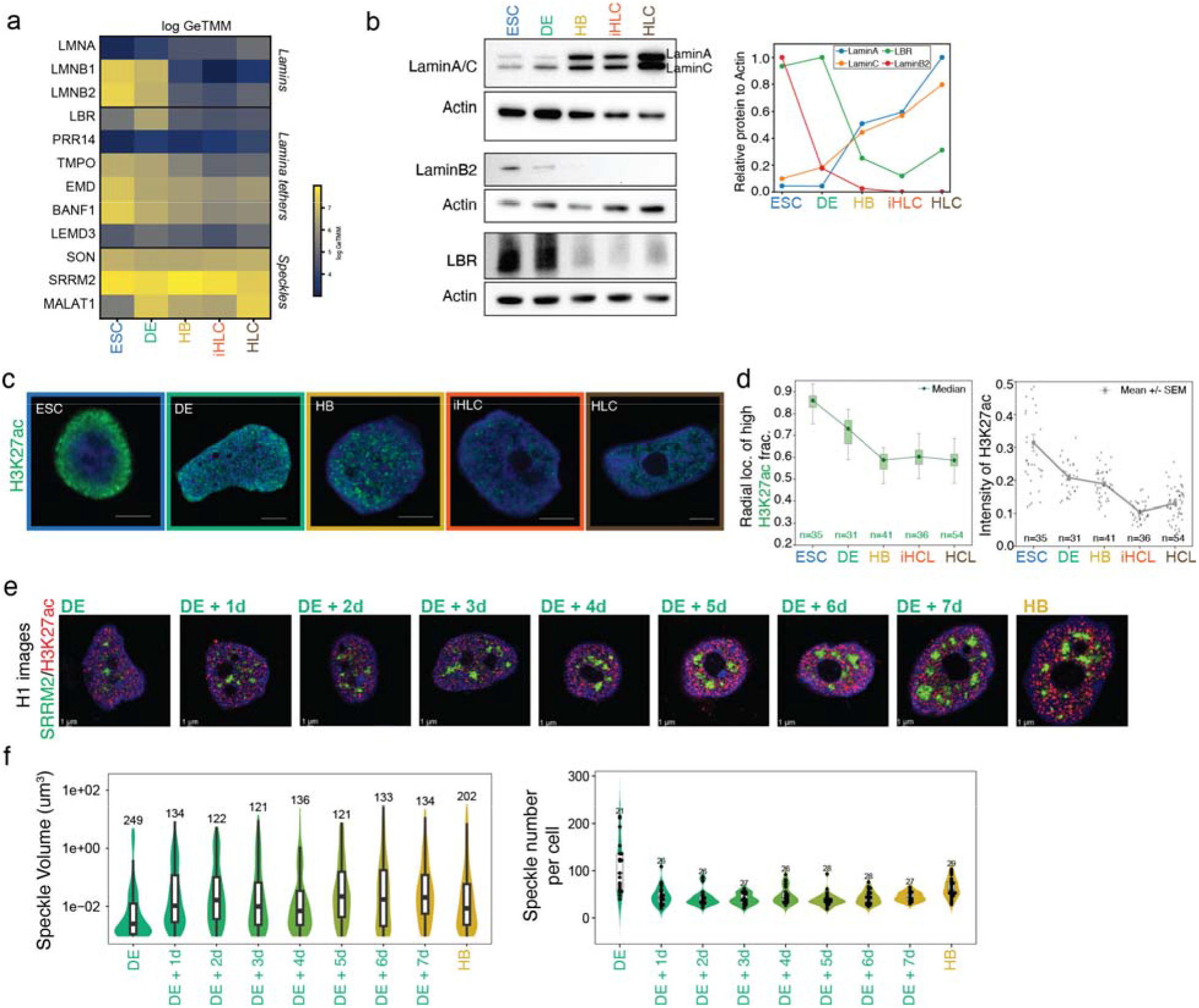
Dynamics of lamin and speckles. **a**, Gene expression level of lamin and speckle components across 5 stages. **b**, Western blots showing protein expression of nuclear lamina components across 5 stages. Quantification of relative protein expression level by normalizing to loading control ACTB (right). **c**, Representative immunofluorescence images of H3K27ac (green, top row) across 5 stages. Scale bar: 5μm. **d**, Quantifications of panel e. Left: Intensity of H3K27ac quantified from H3K27ac immunofluorescence staining across 5 stages. Right: relative radial distance of H3K27ac (green boxplots) high intensity domains in nuclei across 5 stages. **e**, 3D projections of cells stained with SRRM2 during 8-day DE-to-HB transition in H1 cell lines. Scale bar: 5 μm. **f**, Speckle volume (left) and speckle number (right) quantifications for at least 21 cells containing at least 121 speckles for timepoints during 8-day DE-to-HB transition in H1 cell lines (DE = 249, DE + 1d = 134, DE + 2d = 122, DE + 3d = 121, DE + 4d = 136, DE + 5d = 121, DE + 6d = 133, DE + 7d = 134, HB = 202).

**ED Fig. 6:**
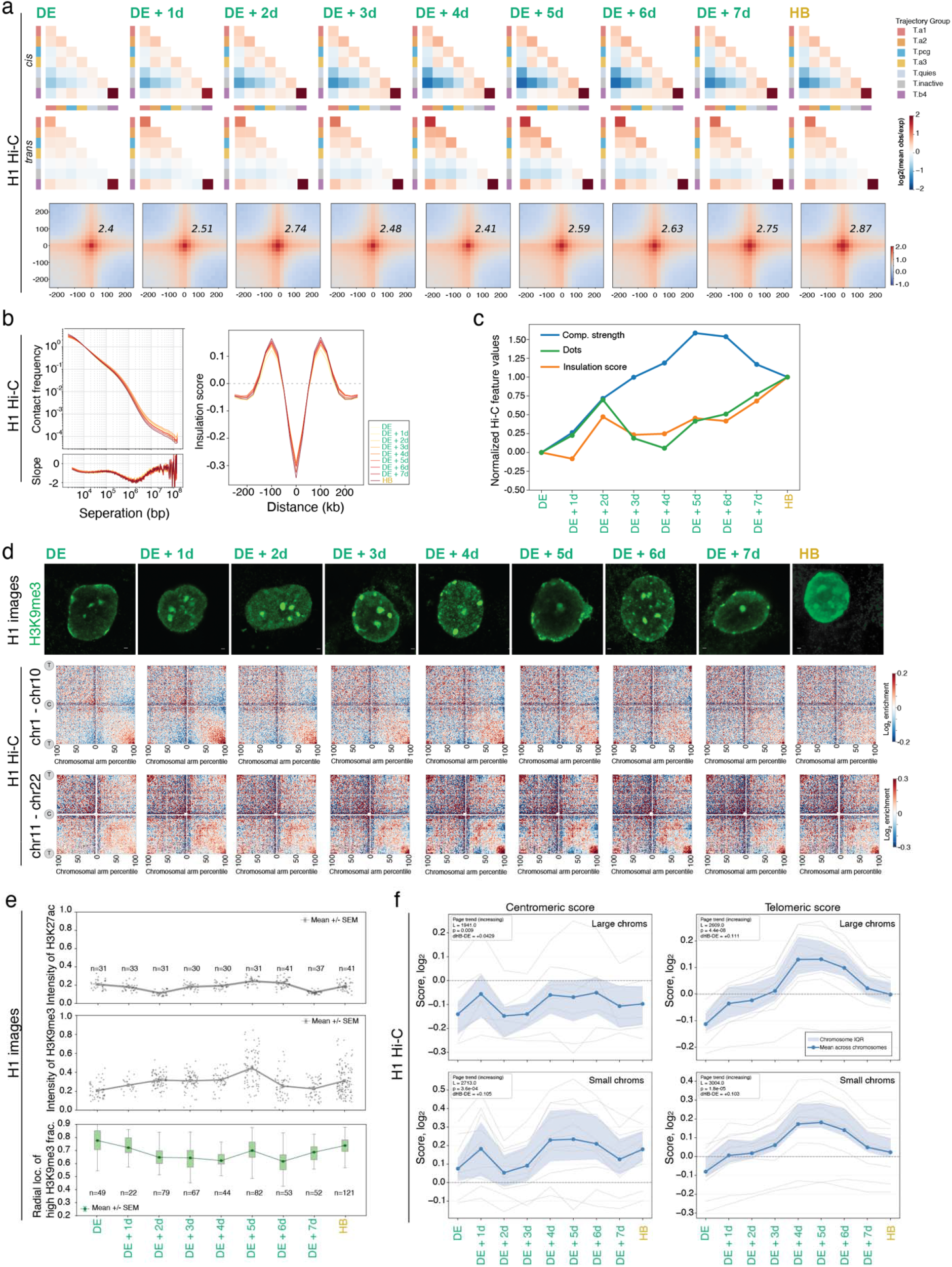
Change of saddle strength, insulation, loops, centromeric and telomeric interactions during the DE to HB transition derived from H1 cells. **a**, Aggregate intrachromosomal (first row) and interchromosomal (second row) contact enrichment between IPT groups across the 8-day DE-to-HB transition. Two-dimensional dot pileups showing associated dot strength across the 8-day DE-to-HB transition (third row, dots calling see methods). **b**, Genome-wide P(s) curves showing average contact frequency at various genomic distances for each stage (top left).and associated slopes (bottom left). Average Hi-C insulation score profiles centered on topological associated domains (TADs) boundaries (right). **c**, Normalized values are scaled from 0 (DE) to 1 (HB) for compartment strength (blue line), insulation score at TAD boundaries (orange line), and loop/dot strength (green line). **d**,Representative immunofluorescence images of H3K9me3 (green) across the 8-day DE-to-HB transition (first row). Scale bar: 10μm. Scaled-chromosome saddle plots of pile-up inter-chromosomal interactions of long chromosomes (chr1-chr10; middle row) or long chromosomes (chr11 - chr22; bottom row) across the 8-day DE-to-HB transition. “T” represents telemeric regions, “C” represents centromeric regions. **e**, Relative average intensity of H3K27ac (first row) and H3K9me3 (middle row) signal in nuclei. Relative radial distance of H3K9me3 high intensity domains in nuclei is also quantified (bottom row). **f**, Quantification of interchromosomal interactions at pericentromeric (right) and telomeric (left) regions across stages. Scores were calculated within 4 Mb of the corresponding telomere/centromere along each chromosome axis. Top panels show interactions among chromosomes 1–10 for pericentromeric (left) and telomeric (right) regions; bottom panels show chromosomes 11–22 for pericentromeric (left) and peri-telomeric (right) regions.

**ED Fig. 7:**
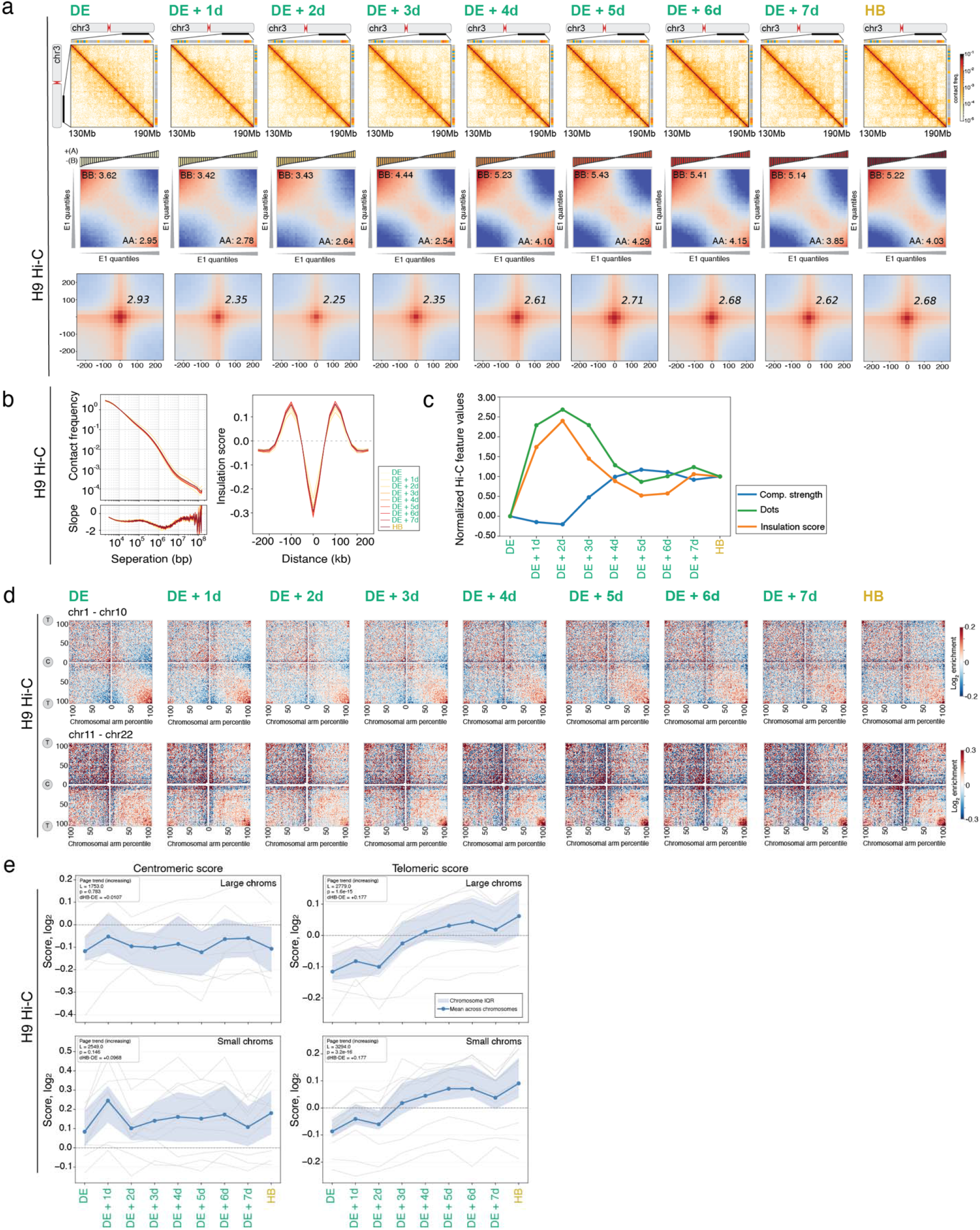
Change in compartmentalization, insulation, loops, centromeric and telomeric interactions during the DE to HB derived from H9 cells. **a**, Representative contact frequency maps for a region on chromosome 3 (chr3: 130Mb-190Mb) across stages (first row). PC1 saddle plots from bulk Hi-C showing compartment strength across stages (middle row). The strength of AA compartment and BB compartments are labeled on the bottom right and the top left of the saddle plot (top 20% bins are used to calculate the strength). Two dimensional dot pileups displaying dot strength for each stage (bottom row, dots calling see methods). **b**, Genome-wide P(s) curves showing average contact frequency at various genomic distances for each stage (top left). and associated slope (bottom left). Average Hi-C insulation score profiles centered on topological associated domains (TADs) boundaries across stages (right). **c**, Normalized values scaled from 0 (DE) to 1 (HB) for compartment strength (blue line), insulation score at TAD boundaries (orange line), and loop/dot strength (green line).**d**, Scaled-chromosome saddle plots of pile-up inter-chromosomal interactions of long chromosomes (chr1-chr10, top row) or long chromosomes (chr11 - chr22; bottom row) across stages. “T” represents telemeric regions, “C” represents centromeric regions. **e**, Quantification of interchromosomal interactions at pericentromeric (right) and telomeric (left) regions across the 8-day DE-to-HB transition in H9 cells. Scores were calculated within 4 Mb of the corresponding telomere/centromere along each chromosome axis. Top panels show interactions among chromosomes 1–10 for pericentromeric (left) and telomeric (right) regions; bottom panels show chromosomes 11–22 for pericentromeric (left) and peritelomeric (right) regions.

**ED Fig. 8:**
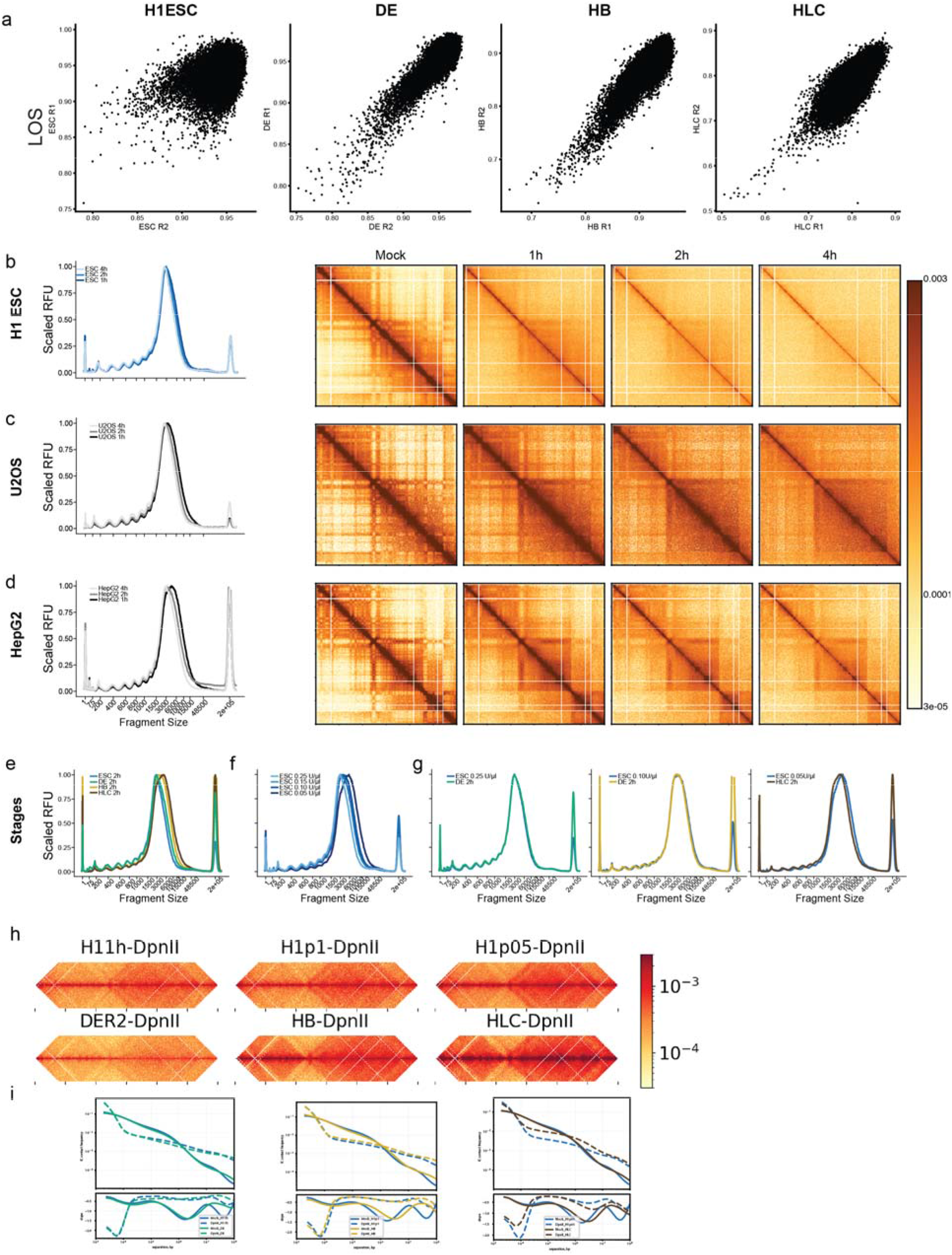
Cell type-specific fragment size distributions and dissolution kinetics. **a**, Scatter plot of LOS values for LC-Hi-C replicates generated in this study. Fragment size analysis and heatmaps resulting from mock-, 1h-, 2h, and 4h DpnII (0.25U/μl) pre-digestion of nuclei isolated from H1 ESCs (**b**), U2OS (**c**) and HepG2 (**d**) cells. **e**, Fragment size distributions generated from 2h DpnII (0.25U/μl) pre-digestion of nuclei isolated from each differentiation stage. **f**, Fragment size distributions generated from pre-digestion of H1 ESC nuclei using various concentrations of DpnII enzyme. **g**, Fragment size distribution matching for ESC/DE, ESC/HB, and ESC, HLC comparisons. **h**, contact frequency heatmaps and scaling (*P(s)*) curve (**i**) comparisons for ESC/DE (left), ESC/HB (middle), and ESC, HLC (right) library pairs that are matched for DpnII fragment size distribution.

